# Cellular gibberellin dynamics govern indeterminate nodule development, morphology and function

**DOI:** 10.1101/2023.09.09.556959

**Authors:** Colleen Drapek, Nadiatul Radzman-Mohd, Annalisa Rizza, Katharina Schiessl, Fabio Dos Santos Barbosa, Jiangqi Wen, Giles E.D. Oldroyd, Alexander M. Jones

## Abstract

During nutrient scarcity, plants can adapt their developmental strategy to maximize their chance of survival. Such plasticity in development is underpinned by hormonal regulation, which mediates the relationship between environmental cues and developmental outputs. In legumes, endosymbiosis with nitrogen fixing bacteria (rhizobia) is a key adaptation for supplying the plant with nitrogen in the form of ammonium. Rhizobia are housed in lateral root-derived organs termed nodules that maintain an environment conducive to Nitrogenase in these bacteria. Several phytohormones are important for regulating the formation of nodules, with both positive and negative roles proposed for gibberellin (GA). In this study, we determined the cellular location and function of bioactive GA during nodule organogenesis using a genetically-encoded second generation GA biosensor, GPS2. We found endogenous bioactive GA accumulates locally at the site of nodule primordia, increasing dramatically in the cortical cell layers, persisting through cell divisions and maintaining accumulation in the mature nodule meristem. We show, through mis-expression of GA catabolic enzymes that suppress GA accumulation, that GA acts as a positive regulator of nodule growth and development. Furthermore, increasing or decreasing GA through perturbation of biosynthesis gene expression can increase or decrease the size of nodules, respectively. This is unique from lateral root formation, a developmental program that shares common organogenesis regulators. We link GA to a wider gene regulatory program by showing that cytokinin as well as nodule-identity genes induce and sustain GA accumulation necessary for nodule function.

## INTRODUCTION

Nutrient acquisition is a fundamental problem facing plants in ecosystems and crops. Fixed nitrogen is particularly scarce in tropical and agricultural soils and its availability is a key determinant for plant health^1^. The nitrogen-fixing clade of plants, which include legumes, have adapted to house nitrogen-fixing bacteria within root lateral organs, termed nodules, as a means of overcoming nitrogen scarcity. Understanding how they do so is important for improving nutrient uptake in commercially important legumes as well as in attempts to improve nitrogen acquisition in cereal crops^2^.

Several phytohormones are important for regulating proper formation and maintenance of nodules. Cytokinin stimulates nodule initiation and accumulates in nodules^3–6^. Its exogenous application results in the formation of spontaneous nodules in several legume species^3–5,7–10^. Cytokinin recruitment of the transcription factor NODULE INCEPTION (NIN) is required for nodule formation and NIN functions in the activation of the key transcriptional gene networks essential for nodule organogenesis and nitrogen fixation^3,7,11–13^. NIN itself is expressed specifically in endosymbiotic conditions, especially in dividing pericycle cells, the epidermis and nodule primordia, which is regulated by both a proximal promoter and distal enhancer elements^14^.

In addition to hormonal cues, transcriptional regulators such as NODULE ROOT 1 and 2 (NOOT1/NOOT2), NF-YA1, LATERAL ORGAN BOUNDARIES DOMAIN proteins 11 and 16 (LBD11/LBD16) and LIGHT SENSITIVE SHORT HYPOCOTYL proteins 1 and 2 (LSH1/2) function in organ development and identity^15–23^. The LSH proteins act downstream of NIN and in roots function specifically in nodule organogenesis. Mutants in *lsh* have deformed, multi-lobed nodules that lack infection and nitrogen-fixation^19^. LSH proteins regulate the NOOTs, mutants of which also lose nodule identity and shape, and ultimately convert to lateral-root like structures^15–17,19^.

Gibberellin (GA) hormones have been increasingly appreciated as having a nuanced but important role in the rhizobia-legume endosymbiosis. DELLA proteins, the antagonists of GA signaling that are degraded in the presence of GA, have been found to be required for rhizobial infection as well as endosymbiotic fungal infection suggesting a negative role for GA in symbiosis^24–26^. In line with this conclusion, treatments with low doses of a GA biosynthesis inhibitor paclobutrazol (PAC) increase nodule number, exogenous GA reduces nodule number and over-expression of nondegradable DELLA can rescue nodulation in cytokinin mutants^25–28^. On the other hand, GA biosynthetic mutants in *Pisum sativum* are deficient in nodulation^29,30^, GA biosynthesis genes in *Lotus japonicus* and *Glycine max* are expressed at the site of nodule organogenesis, and there is a general trend of upregulation of GA-biosynthetic genes across transcriptomes of nodulation^18,31–38^. Recently, *Medicago truncatula* mutants deficient in the final stage of GA biosynthesis were found to have fewer nodules than wild-type counterparts^36^. Further, *Agrobacterium rhizogenes-*mediated transient expression of Gibberellin Perception Sensor 1, a FRET biosensor for GA, in soybean hairy roots showed possible accumulation in nodule primordia cells, in agreement with LC-MS detection of GA accumulation in nodules^34,35^. Rhizobia themselves can also produce GA, which may give them a fitness advantage post-nodule senescence.^39^

These studies of GA function in nodulation imply both positive and negative effects, suggesting finely-tuned control of endogenous GA levels or their distribution in space or time. Here, we use an improved FRET biosensor for GA, nlsGPS2 (Griffiths et. al. 2023), genetically encoded in *M. truncatula,* to visualize cellular GA dynamics *in vivo* and address where and when GA is important for nodule development and maintenance. In early nodule development, we see a striking pattern of GA accumulation in dividing cortical cells of nodule primordia that is maintained in the mature nodule apex. We determine GA is key in the early initiation of cell division leading to the nodule primordia, size and function. This is in stark contrast to the initiation of lateral roots, that show little to no GA accumulation during organogenesis. Symbiotic mutants that lose their organ identity and convert to root-like structures lose their GA accumulation pattern. We conclude that GA is a key differentiator between nodule and lateral root identity.

## RESULTS

### A genetically-encoded second generation GA sensor shows a robust pattern of GA accumulation during nodule organogenesis

In order to understand the time, place and levels of cellular GA in the rhizobia-legume symbiosis, we stably transformed *M. truncatula* with a second generation GA biosensor, nlsGPS2, under control of the *L. japonicus* ubiquitin promoter. Use of a biosensor permits visualization of bioactive GA at cell resolution, and thus offers spatial resolution where other techniques, such as chromatography/mass spectrometry, quantify but do not spatially resolve hormone species. The nlsGPS2 sensor was optimized for use in *A. thaliana* ^40,41^, thus we confirmed the functionality of the sensor in a new species by quantifying emission ratio changes in roots treated with exogenous bioactive GA species (GA_4_ and GA_3_) in comparison to mock treatments (Fig. S1).

We then examined nodule development from initiation to maturation over a series of days after inoculation with *Sinorhizobium meliloti* strain 2011 (Sm2011). Prior to infection, GA is low in *M. truncatula* roots (Fig. 1a), but at 4 days post infection (dpi), GA accumulates in dividing cortical cells with low levels in the stele, endodermis and epidermis as observed in radial sections (Fig. 1b). In 5 dpi whole mounted roots, there is a robust pattern of a high GA gradient decreasing from the centre to the periphery of the nodule primordium (Fig. 1c,e). The GA remains low in the root surrounding the developing nodule, an intriguing result given the root-wide expression of GA biosynthesis genes during infection^36^ (Fig. 1c). At maturity, GA accumulation remains high towards the nodule apex, but low in the surrounding tissue and stele (Fig. 1e,f), a striking difference from reports in soybean (determinate) nodules, where GA was reported to diminish at the nodule apex after several days^35^. Note that we do not see expression of the sensor in the central zone and suggest this is due to the tightly-regulated oxygen environment of infected cells (see discussion).

**Fig.1.**
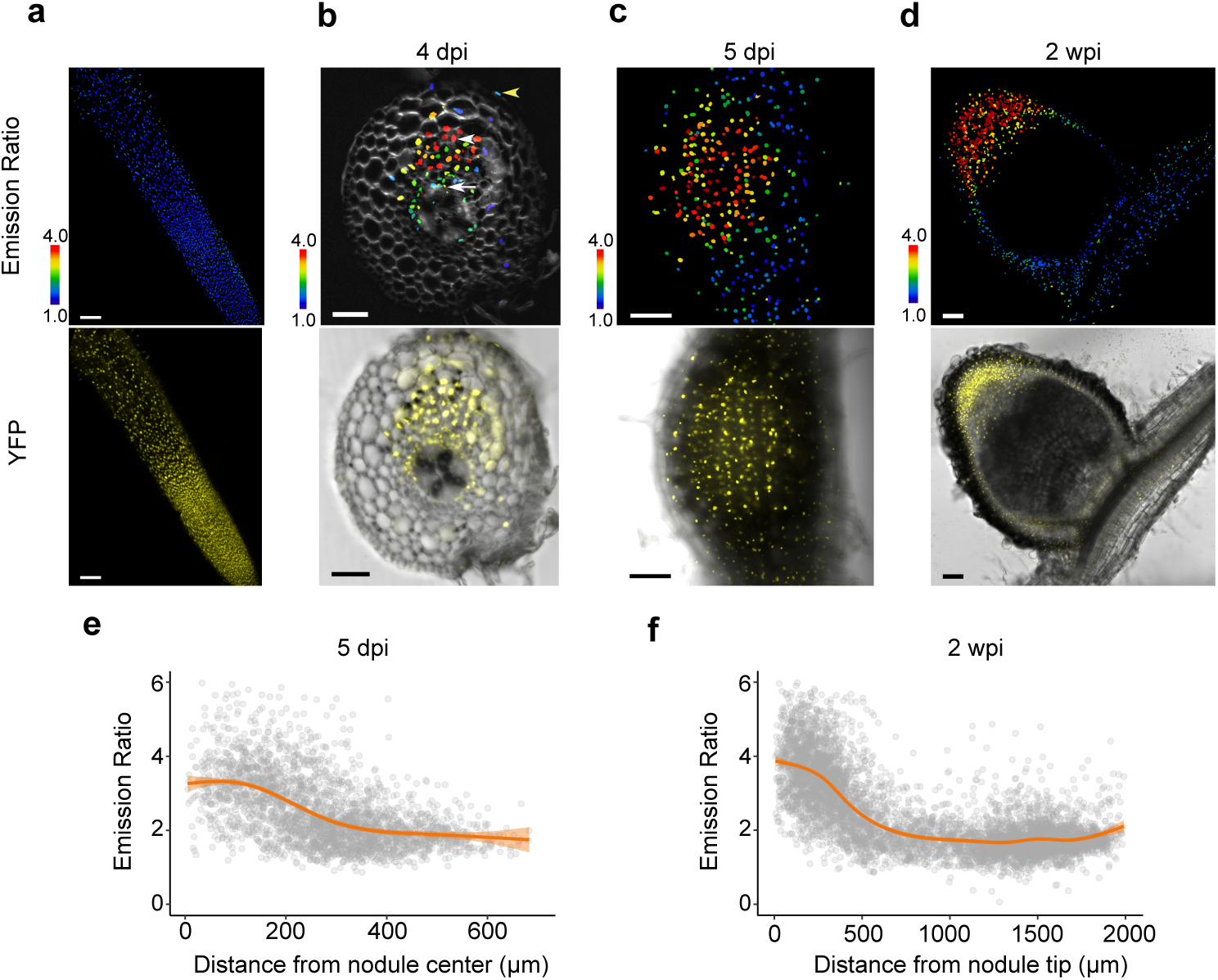
GA accumulates early in nodule development and persists in the nodule apex. **(a)** Emission ratio of *LjUBQp::*nlsGPS2 in *M. truncatula* root (top panel) and YFP control (bottom panel), N ≥ 20. **(b-d)** Emission ratio of nlsGPS2 and YFP/Brightfield channel overlay in *M. truncatula* nodules inoculated with *Sm2011*. **(b)** 4 dpi nodule primordia embedded in 4.5% agarose and sliced in 100µm sections, N=3. White arrowhead indicates cortex, white arrow indicates endodermis/stele, yellow arrowhead indicates epidermis. **(c)** 5 dpi whole mount nodule primordia, N ≥ 20. **(d)** 2 wpi nodule embedded 4.5% agarose and sliced in 100µm sections, N ≥ 20. **(e)** Emission ratio of nuclei (individual dots) from whole mount nodules at 5 dpi as a function of distance from nodule center, N=9. Curves of best fit are computed in R using a generalized additive model via ggplot. **(f)** Quantification of emission ratio of nuclei (individual dots) of 2 wpi nodules as a function of distance from nodule tip, N=5. Curves of best fit are computed in R using a generalized additive model via ggplot. Bars = 100 µm in (**a,c,d**); Bar = 50µm in (**b**). Data shown are from >3 biological replicates. Additional biological replicates, CFP and FRET channels available in Fig. S2.

### The mechanism underlying spatial restriction of GA

Given the strikingly tight localization of GA accumulation during nodule primordia formation and at maturity, we sought to determine how the spatial accumulation of GA is formed (Fig. 2a). Gibberellin 20-oxidase (GA20ox), which catalyzes the penultimate step in GA biosynthesis, has traditionally been seen as the rate limiting enzyme in bioactive GA production. However, this does not capture the enzymatic biology relevant in all scenarios (for example, GA3ox that catalyzes the last biosynthetic step, but not GA20ox, is rate limiting in the Arabidopsis root elongation zone)^41^. We therefore supplied *M. truncatula* roots with the precursors for GA20ox (GA_12_) and GA3ox (GA_9_) to test if either early biosynthetic steps or GA20ox activity were rate limiting for GA accumulation and responsible for GA distribution in nodule development (Fig. 2b-e). Treatment with GA_12_ at the time of infection quantifiably increased GA within the nodule primordia, but did not alter the generally centralized accumulation of GA (Fig. 2c). Similarly, treatment with GA_9_ quantifiably increased GA levels around the nodule but did not abolish the general pattern of central GA accumulation, indicating that while prior enzymatic steps quantitatively contribute to setting GA levels in nodulation, limitation of GA3ox activity or GA depletion mechanisms exert important control over GA levels and distribution (Fig. 2d). Treatment with a bioactive GA, GA_3_, flattens the accumulation across the entire root (Fig. 2e,g). Previously, it was shown that soybean has a nodule-specific GA20ox, and that MtNIN can activate expression of MtGA3ox1 broadly in *M. truncatula*^35,36^. We examined *A. rhizogenes* transformed roots containing the 3 kb upstream region of MtGA20ox1 (Medtr1g102070) and MtGA3ox1 (Medtr2g102570). In agreement with previously published transcriptome and promoter studies, both genes are broadly expressed in the root (Fig. S3). We conclude that the pattern of GA accumulation during nodule development is not defined solely by the expression of the rate-limiting enzymes involved in GA biosynthesis, which are broadly expressed, but rather must also involve post-transcriptional regulation of GA3ox activity or GA depletion (i.e. export and catabolism).

**Fig. 2.**
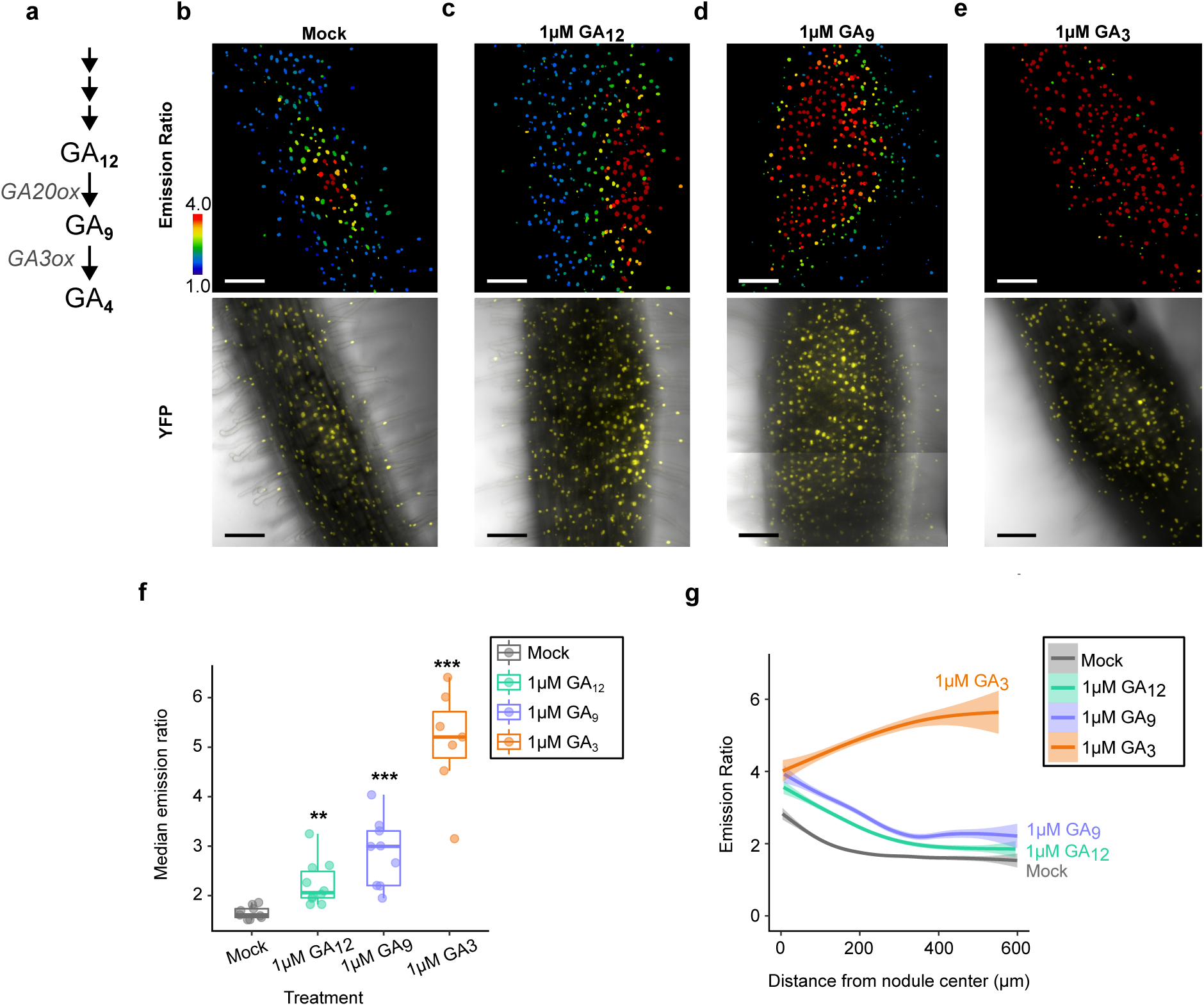
Spatial limitation of GA partially depends on precursor availability. **(a)** A simplified schematic of the final metabolic steps in bioactive GA biosynthesis. Enzyme families are italicized. **(b-e)** Emission ratio and YFP/Brightfield overlay of nlsGPS2 in developing nodules (5 dpi) treated with **(b)** mock **(c)** 1µM GA12 **(d)** 1 µM GA9 and **(e)** 1 µM GA3, a bioactive GA mimic. Treatment was carried out at time of infection. **(f)** Emission ratio by treatment. Each dot represents median emission ratio per sample. Welch’s t-test between treatment and mock **p-value **<0.01, ***<0.001. Median values: mock = 1.63, 1 µM GA_12_ = 2.08, 1µM GA_9_ = 3.01, 1µM GA_3_ = 5.22. N≥7 from three biological replicates **(g)** Emission ratio of nuclei relative by treatment as a function of distance from nodule center. Curves of best fit are computed in R using a generalized additive model via ggplot. N≥7. Bars = 100 µm.

Rhizobia themselves can produce bioactive GA and GA precursors that affect nodule size^39^. We asked if the initial source of the GA detected with nlsGPS2 could be from rhizobia. To test this, we asked if spontaneous nodules that lack rhizobia contained bioactive GA. We induced spontaneous nodules by introducing a ubiquitously-expressed, constitutively active CCaMK into the nlsGPS2 lines via *A. rhizogenes* transformation^42^. These nodules show a GA accumulation pattern similar to infected plants (Fig. S4), indicating that, at least at nodule onset, GA is produced by the host.

### GA is required in cortical cells to govern nodule development and size

Given the pattern of GA accumulation in nodule primordia, we explored if the accumulation of GA in nodules is functionally important. To do this, we sought to disrupt early accumulation of GA in nodules. We treated plants with 0.1 µM and 1 µM PAC at the time of infection, which we found, in agreement with previously published reports, resulted in more nodules per plant^25,26^. However, when we examined these nodules for GA accumulation, neither concentration resulted in a lack of GA in developing nodules, though there was a quantifiable decrease in the overall distribution (Fig. 3a-b). Therefore, the increase in nodulation in PAC treatment is not due to a lack of GA in developing nodules, and is consistent with a role for DELLA, and therefore GA depletion, elsewhere during initial infection.

**Fig. 3.**
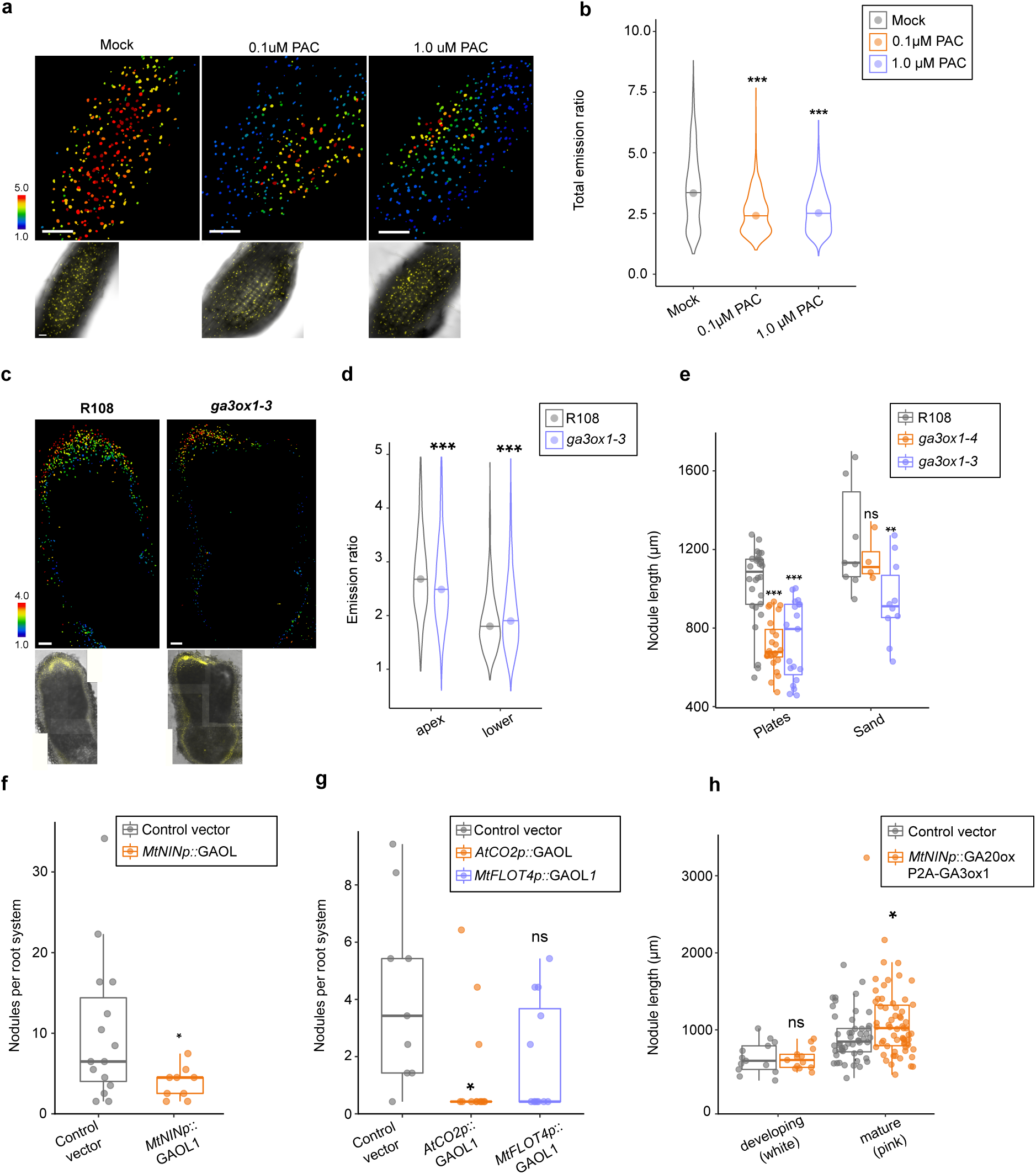
GA functions in early nodule development in cortical cells. **(a)** Emission ratio (above) and Brightfield/YFP insets (below) of nlsGPS2 in mock and PAC-treated 5 dpi developing nodules. Treatment was carried out at time of inoculation. **(b)** Distribution of emission ratio of nuclei in mock, 0.1µM PAC and 1.0µM PAC-treated samples. Mock quantification data is also represented in Fig. 2f-g. Welch’s t-test between mock and indicated treatment, ***p-value<0.001. Median values: mock = 3.3445, 0.1 µM PAC = 2.4125, 1.0 µM PAC = 2.5195. N≥3. Representative replicate from three biological replicates is shown. **(c)** Emission ratio (above) and Brightfield/YFP insets (below) of nlsGPS2 in wild-type and *ga3ox1-3* nodules 4 wpi in sand mix. **(d)** Distribution of emission ratio of nuclei in the nodule apex and in the lower nodule/root in R108 compared to *ga3ox1-3* in 4 wpi in sand mix. N≥5 nodules, Welch’s test *\*\*\**p-value < 0.001. Median values apex: R108 = 2.736, *ga3ox1-3* = 2.56. Median values lower: R108 = 1.851, *ga3ox1-3* = 1.956. Representative replicate from three biological replicates is shown. **(e)** Nodule length from base to tip at 2 wpi in R108, *ga3ox1*-3 and *ga3ox1-4* on plates or in sand. In the plates assay, Welch’s t-test *** p-value <0.001; in the sand assay, Student’s t-test **p-value <0.01. Median values plates: ; median values sand: .**(f)** Total nodules per root system in *A.rhizogenes*-transformed roots containing a control construct (*AtUBQp::*AtPIP2A-mScarlet) or the GA-catabolic enzyme *MtGAOL1* under control of a proximal *NIN* promoter containing distal elements. Wilcoxon rank sum test *p-value < 0.05; N≥10 root systems. Median values: control = 5, *MtNINp::*MtGAOL1 = 3.**(g)** Total nodules per root system in *A.rhizogenes*-transformed roots containing a control construct (*AtUBQp::* AtPIP2A-mScarlet), the GA-catabolic enzyme *MtGAOL1* under control of the cortex-specific *A. thaliana* promoter *AtCO2* or epidermis-specific promoter *MtFLOT4*. Wilcox rank sum test **p-value <0.01, N≥10 root systems. Median values: control = 3, *AtCO2p*::MtGAOL1 = 0, *MtFLOT4p*::MtGAOL1 = 0. **(h)** Nodule length (µm) from base to tip of *A.rhizogenes-*transformed roots containing a control construct (*AtUBQp::* AtPIP2A-mScarlet) or increased local GA biosynthesis (*MtNINp::*GA20ox1-P2A-GA3ox1) in developing (white) nodules and nitrogen-fixing (pink) nodules. Welch’s t-test *p-value <0.05. Median values immature (white): control = 599.687, *MtNINp::*GA20ox1-P2A-GA3ox1 = 609.544; Median values mature (pink): control = 849.228, *MtNINp::*GA20ox1-P2A-GA3ox1 = 1023.115. N≥50. Additional *A. rhizogenes* replicates displayed in Fig S6.

We therefore explored a genetic approach to deplete GA using mutants for *MtGA3ox1,* that is highly expressed during the rhizobial symbiosis^18,31–33,36–38^. We selected two insertional alleles for MtGA3ox1, NF19087 (*ga3ox1-3*) and NF13294 (*ga3ox1-4*). Both are characteristically dwarfed, slow to germinate, and had short internodes, consistent with GA-related phenotypes^43^. Despite these strong GA-related phenotypes, we observed only subtle effects in nodulation (Fig. S5), counter to what has been previously reported for different alleles^36^. We examined nlsGPS2 in the *ga3ox1-3* background, and found GA was still present, indicating this mutant cannot entirely remove GA from the nodule, pointing at likely compensatory genes, for instance a second copy of GA3ox, *MtGA3ox2* (Medtr1g011580) (Fig. 3c-d). While the phenotypes were more subtle than previously reported, we did observe a reduction in nodule size (Fig. 3e), consistent with reports for biosynthetic mutants in pea^29^. Symbiosomes appeared normal in nodules of *ga3ox1* (Fig. S5).

To better control GA depletion, we mis-expressed the *GA2-oxidase like (GAOL)* gene *MtGAOL1* (Medtr3g464530), an enzyme in the *Elongated Uppermost Internode-like* family of cytochrome P450 monooxygenases that breaks down bioactive GA. A similar approach using GA catabolic enzymes has been used in *A. thaliana* for spatially restricting GA^44^. Misexpression of *MtGAOL1,* under a fully functional *NIN* promoter composed of distal and proximal regions of the promoter^14^, led to a marked reduction in the total number of nodules and the absence of pink nodules (indicative of the presence of leghemoglobin for functional nitrogen fixation; Fig. 3f, Fig. S6). We also expressed *MtGAOL1* under cortical-specific (*A. thaliana CO2*) and epidermal-specific FLOTILLIN4 (*MtFLOT4*) promoters^28,45–47^. Cortical expression of *MtGAOL1* resulted in a severe reduction in developing nodules, whereas epidermal expression of *MtGAOL1* had a trend in nodule reduction but no significant effect (Fig. 3g, Fig. S6). Together, these results indicate a role for symbiotic GA accumulation in the behavior of root cortical cells leading to nodule development. To further explore this role, we attempted to locally increase GA levels in the nodule primordia. We expressed both *MtGA20ox1* and *MtGA3ox1* together, under the *NIN* promoter, in *A. rhizogenes* transformed *M. truncatula* roots. These two enzymes are key steps in GA biosynthesis^48^ and consistently, we observed quantifiably higher levels of GA in these nodules (Fig. S7). While nodule number did not increase in these roots (Fig. S7), mature nodule size was significantly larger in mature nodules (Fig. 3h). We conclude that GA is a positive regulator during nodule organogenesis initiation and development.

### GA accumulates downstream of cytokinin in a *NIN*-independent manner

Cytokinin is a major regulator of nodule organogenesis that acts upstream of the transcription factor *NIN*^3^. In many legume species, exogenous cytokinin treatment can induce spontaneous nodule formation and root cortical cell divisions^10^. Transcriptomic data show evidence that cytokinin signalling may be the principle regulator of GA biosynthesis during nodulation given that all genes in the GA biosynthesis pathway that are significantly up or down-regulated during rhizobia inoculation are absent in the cytokinin receptor mutant *cre1*^18^. We therefore hypothesized GA accumulation acts downstream of cytokinin and consistently treatment with 6-benzylamino-purine (BA) for 24h results in the accumulation of GA in *M. truncatula* roots (Fig 4a). More recently, it was shown in *M. truncatula* that the broadly-expressed upregulation of *MtGA3ox1* during symbiosis is dependent on NIN^36^ and therefore we tested if BA-induced GA accumulation in the *nin-4* mutant. Surprisingly, we observed that BA-induced GA accumulation is independent of *NIN* (Fig. 4a-b) but that rhizobia-induced GA accumulation is *NIN*-dependent (Fig. 4c-d). These results imply a coherent feedforward loop with cytokinin recruiting NIN for stimulating GA accumulation in a symbiotic context.

**Fig. 4.**
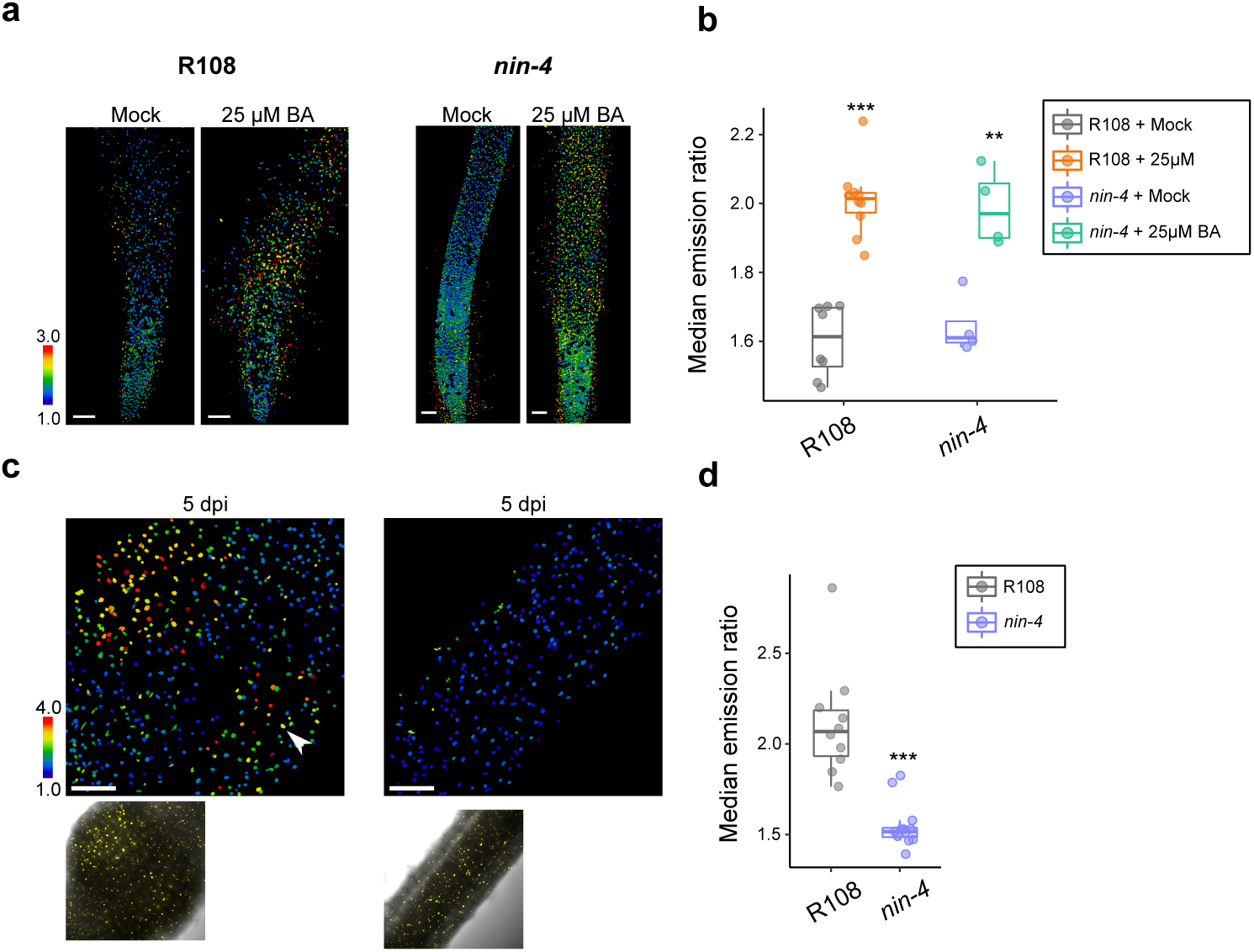
Cytokinin can recruit GA accumulation in a NIN-independent fashion. **(a)** Emission ratio of nlsGPS2 in either R108 or *nin-4* mock treated compared roots compared to 24h 25µM BA-treated roots. Samples are 6 days old. **(b)** Emission ratio from indicated genotypes and treatments. Each dot represents median emission ratio from one sample. Wilcoxon rank sum test **p-value < 0.01, ***p-value < 0.001. Median values: R108 mock = 1.613, R108 treated = 2.014; *nin-4* mock = 1.61, *nin-4* treated = 1.970. **(c)** Emission ratio of nlsGPS2 (top panel) and YFP/brightfield overlay (bottom panel) in R108 (left) and *nin-4* (right) at 5 dpi. Images taken at point of susceptibility zone at time infection. Arrowhead indicates second nodule forming. **(d)** Median emission ratio from indicated genotype. Each dot represents one sample. Wilcoxon rank sum test ***p-value < 0.001. Median values: R108 = 2.069, *nin-4* = 1.517. **(b,d)** N>=10. Data shown are from three biological replicates.

### GA accumulation is a signature of nodule maturation and is regulated by organ-identity transcription factors

We noticed lateral roots failed to accumulate GA (Fig. 5a). This is an intriguing difference as there is overlap in the developmental organogenesis programs of lateral roots and nodules^18,20^.This suggests that GA accumulation acts as an important differentiator between nodule and lateral root development, and as such may be controlled by nodule-specific identity regulators, *NOOT* and *LSH* ^15–17,19^ We introduced the nlsGPS2 sensor into *noot* and *lsh* and found disturbances in the GA gradients (Fig. 5b-f). Consistent with reported phenotypes, there is a range of nodule structures that occur in these mutant backgrounds and we sought to categorize them by (1) nodule region (nodule apex, lower nodule, or converted root) and (2) degree of organ conversion. For degree of organ conversion, we considered mutant organs as type I if they maintained rounded morphology, type II if they had both a rounded base and lateral-root like structure and type III if they were fully converted with a centralized vasculature similar to lateral roots (only found in *noot1noot2*). In *lsh1lsh2* mutants, GA is significantly lower in the nodule apex compared to wildtype plants, and significantly higher in surrounding tissue, implying a general inability to localize GA accumulation (Fig. 5b-c). Overall GA levels in *lsh1lsh2* mutants were not significantly different in type I organs but are significantly lower in converted root-like structures (Fig. 5b-d). Similar to *lsh1lsh2*, there was significantly less GA in nodule apices of *noot1/noot2* mutants and overall GA was significantly lower in converted organ structures (Fig. 5e-g). These results underpin the observation that GA accumulation strongly correlates with root organ identity, in that as nodules in these mutants convert to lateral roots, we observe an abolition of GA accumulation.

**Fig. 5.**
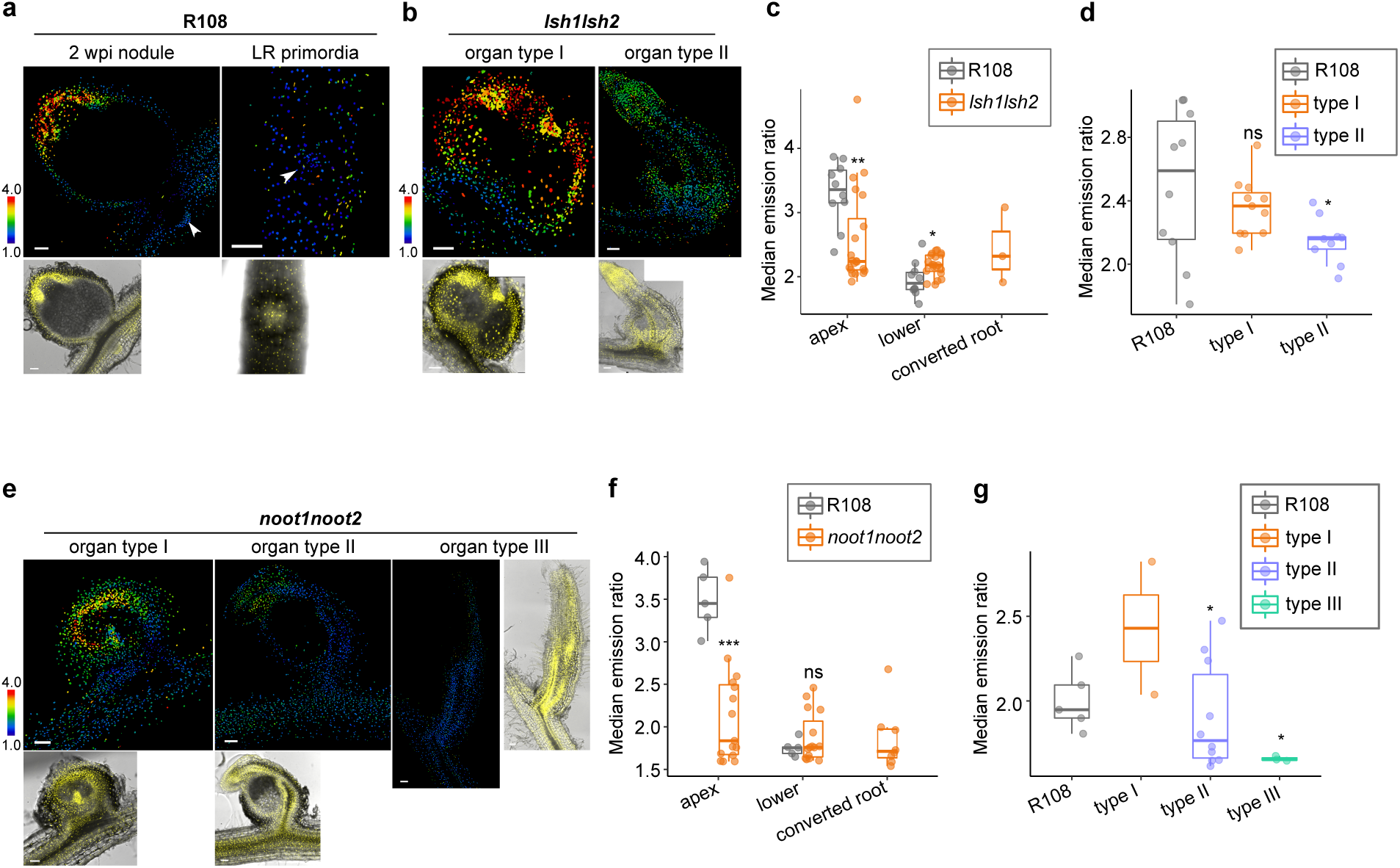
Transcription factors *LSH1LSH2* and *NOOT1NOOT2* regulate and maintain GA gradients in mature nodules. (**a**) Emission ratio of nlsGPS2 (top panel) and YFP/Brightfield (bottom panel) of an R108 2 wpi nodule embedded in 4.5% agarose and sliced in 100 µm sections (left) and lateral root (LR) primordia (right) from a 5 day old sample. Arrowheads indicate LR primordia. (**b**) Emission ratio of nlsGPS2 (top panel) and YFP/brightfield (bottom panel) of a 2 wpi *lsh1lsh2* nodule embedded in 4.5% agarose and sliced. Left is type I (round) organ and right is type II (converting) organ. (**c**) Median emission ratio of nodule regions from indicated genotypes. For apex, wilcoxon rank sum test, **p-value <0.01, median values: R108 = 3.355; *lsh1lsh2* = 2.23775. For lower, Student’s t-test *p-value<0.05, median values R108 = 1.897, *lsh1lsh2* = 2.1875. N≥9 from three biological replicates. **(d)** Median emission ratio of entire nodule region from indicated organ type. Welch’s t-test *p-value <0.05, median values: R108 = 2.589, type I = 2.367, type II = 2.16. N≥9 from three biological replicates. **(e)** Emission ratio of nlsGPS2 (top panel) and YFP/brightfield (bottom/right panel) of a 2 wpi *noot1noot2* nodule embedded in 4.5% agarose and sliced 100 µm sections. Left is type I (round) organ, center is type II (converting) organ, and right is type II (fully converted) organ. **(f)** Median emission ratio of nodule regions from indicated genotypes. For apex, Wilcoxon rank sum test ***p-value<0.001, median values: R108 = 3.45, *noot1noot2* = 1.836. For lower, median values R108 = 1.755, *noot1noot2* = 1.759. N≥5. (d). Welch’s T-test, ***p-value <0.001, N≥5. **(g)** Median emission ratio of entire nodule region from indicated organ type. For type II, Wilcoxon rank sum test *p-value<0.05; median values: R108 = 1.948, *noot1noot2* = 1.766, N≥5. For type III, Welch’s t-test *p-value<0.05, median values: R108 = 1.948, *noot1noot2* = 1.653. N≥3. In **(a,b, d)** Bar = 100µm. Data shown are from three biological replicates.

We sought to determine if supplementing GA could help rescue either organ morphology or nitrogen fixation. We treated *noot* mutants with 100nM GA_4_ at 4 dpi for a further 10 days (for a total of 14 dpi), so as to supply GA only after the initiation of rhizobial infection, a process highly dependent on the action of DELLAs^25,26^. We found no obvious changes in nitrogen fixation or organ conversion, but found post-initiation GA treatment was sufficient to rescue nodule size in *lsh1lsh2* mutants to wildtype, a further link between GA levels and nodule size (Fig. S8). This implies that, while *LSH* and *NOOT* have a role in regulating GA distribution, they have other functions in organ identity maintenance that cannot be solely explained by loss of GA accumulation and that GA loss does not alone explain organ conversion.

### GA accumulation functions in maintaining mature nodule morphology and function

We observed GA accumulation at both early initiation, but also in mature nodules, suggesting a possible function for GA throughout nodule development. To test for functions at the later stages of nodule development, we treated roots with high doses of PAC (20µM) at 7 days after initial infection, and for a further 7 days, so as to not to affect infection or early nodule development. We found treatment with 20µM PAC significantly decreased GA levels in the mature nodule apex and increased GA levels in the region outside of the nodule (Fig. 6e-g). These roots had a significant decrease in mature nodules, an increase in immature white nodules, and a decrease in overall nodule size (Fig. 6a-d). Taken together, this demonstrates the GA distribution of high in the apex and lower in surrounding tissues functions in post-nodule initiation stages to promote nodule growth and maturation to nitrogen-fixation.

**Fig. 6.**
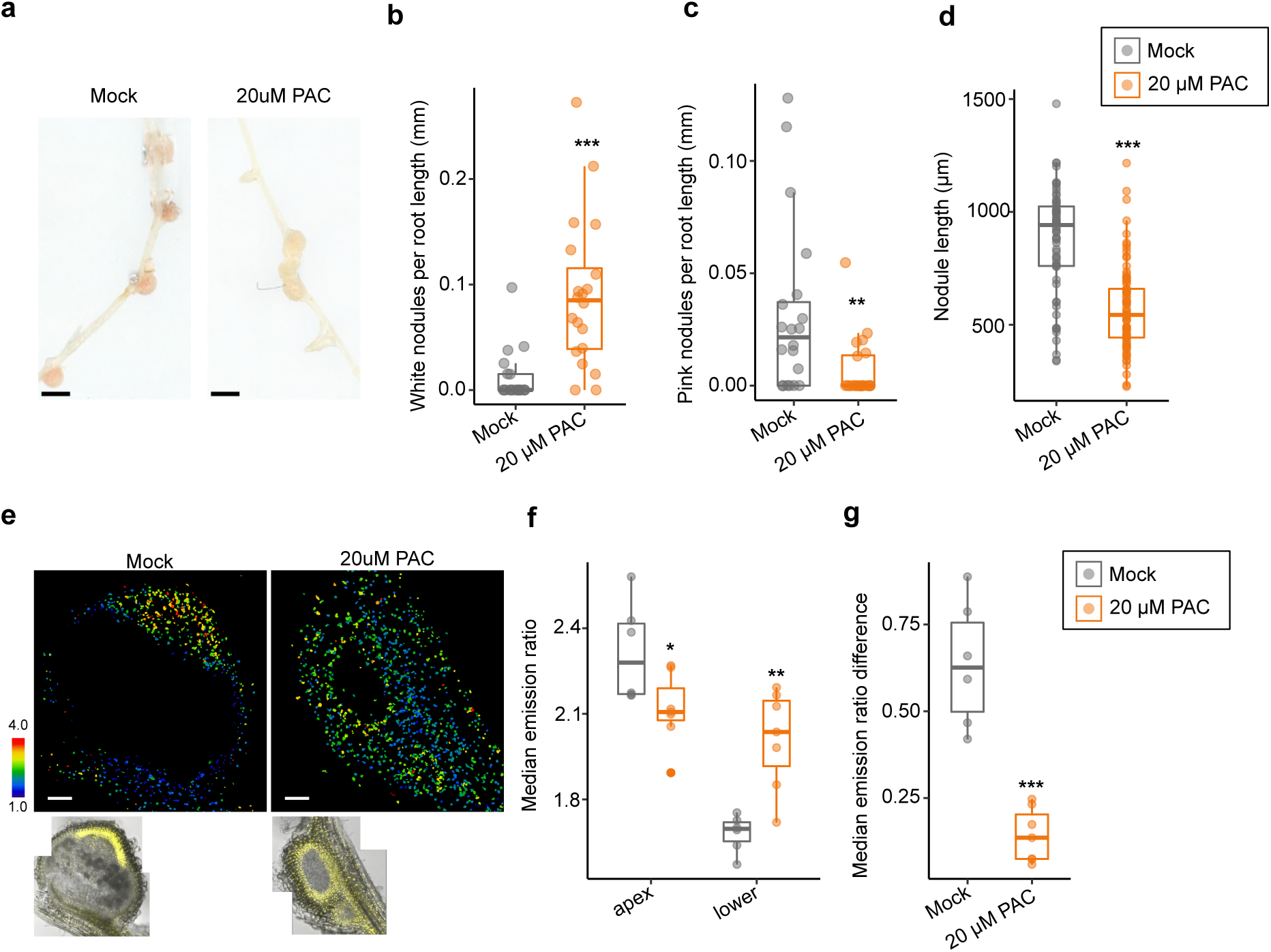
GA patterning is required to maintain nodule growth and nitrogen fixation in mature nodules. **(a)** Scanned representative images of 14 dpi nodules from mock treated roots (left) and roots treated with 20µM PAC for 7 days at 7dpi. Bar = 1mm. **(b)** Quantification of average white nodules per root length (mm). Welch’s t-test ***p-value < 0.001, median values mock = 0, PAC =0.0849. N=20 (**c**) Quantification of average pink nodules per root length (mm). Wilcoxon rank sum test **p-value < 0.01, median values: mock = 0.0215; 20 µM PAC = 0. **(d)** Quantification of nodule length (µm) from mock-treated and 20µM PAC-treated roots. Wlech’s t-test ***p-value < 0.001, median values: mock = 941.5, 20µM PAC = 544. N ≥ 62. **(e)** Emission ratio images (top panel) and YFP/Brightfield controls (bottom panel) of a mock-treated nodule and a nodule from roots treated for 7 days with 20µM PAC. **(f)** Emission ratio from nodule apex region or lower nodule/root. Each dot is median emission from an individual. Student’s t-test *p-value < 0.05, **p-value < 0.01. Median apex values: mock = 2.294, 20µM PAC = 2.12. Median lower values: mock = 1.711, 20µM PAC = 2.051. N≥ 5. **(g)** Absolute value of the median emission ratio difference between the apex and lower region of a nodule. Student’s t-test ***p-value < 0.001. N≥ 5. Data shown are from three biological replicates.

## DISCUSSION

The studies of GA function in nodulation in multiple species of legumes present a varied and sometimes confounding picture. Nonetheless, it was clear that GA accumulates during the association with rhizobia and that the DELLA proteins, which are negatively regulated by GA, are positive activators of rhizobial infection. In this study we use a high-resolution GA FRET biosensor, alongside targeted genetic manipulation, to demonstrate that GA acts as an essential positive regulator of early cortical cell division and differentiation required for nodule initiation as well as later nodule growth and maturation. GA accumulation is closely correlated with nodule-identity: GA accumulation is a strong differentiator between nodule and lateral root development and as nodules lose their specific identity, it correlates with a loss of GA accumulation. As such, GA accumulation has opposing functions in the process of nodulation, a positive regulator of cell divisions leading to nodule meristem formation, while also antagonizing rhizobial infection. This contrasting functionality in organogenesis versus infection helps explain the sometimes contradictory results and conclusions of previous studies.

Central to this study is the stable introduction of a GA biosensor into *M. truncatula,* that for the first time allows an assessment of cellular GA levels throughout the process of nodulation. Our results indicate that GA accumulates early during the initiation of cell division in cortical cells, but remains at relatively low levels in the outer cortex and epidermis where rhizobial infection is initiated (Fig. 1b). Such a spatial separation of GA accumulation would allow DELLA accumulation in outer root layers, facilitating rhizobial infection, while at the same time promoting cell divisions in the root cortex. By technical limitation, we do not see the sensor expressed in the central zone of the mature nodule. We theorize this is due to the tight regulation of oxygen in these cells, which is limited for the oxygen-intolerant nitrogenase enzyme^49^. The GFP-derived fluorophores of nlsGPS2 require oxygen for their maturation^50^, which may not be readily available in these cells.

GA accumulation in dividing, cortical cells of the nodule primordia is functionally important for promoting nodule growth and development. This is an unusual example of GA association with dividing cells, suggesting atypical meristem gene regulation in indeterminate nodules. Typically, GA has been linked to cell elongation and cell growth rather than division, for example in the elongation of hypocotyl cells and in shoot internodes^51,52^. However, *in vivo* analysis of GA biosynthesis in *Arabidopsis* primary root tips expressing nlsGPS1 revealed distinct enzyme regulatory regimes for apical cells of the division zone versus elongation zone cells^41^. Furthermore, targeted manipulation of GA levels in the same tissues indicated that GA promotes division zone length and cell elongation, likely via spatially distinct accumulations^44^. Lateral meristem identity has also been found to be controlled by GA in grapevine tendrils and strawberry runners, providing useful examples for comparing to the patterns in the nodule apex^53–55^.

We show the pattern of GA accumulation in nodules is important for governing size and function (Fig. 6), and patterning is downstream of several key components involved in nodule initiation and identity. While bioactive GA accumulation is limited by precursor availability (Fig. 2e-f), spatial restriction of GA does not solely depend on it. This suggests additional layers of tissue regulation, such as localized biosynthetic enzymatic activity, transport or catabolism. Consistent with this, the catabolic gene *MtGA2ox10* gene (MTR) has been found to play a role in nodule development and infection thread formation^56^ and *MtGA2ox2*, another catabolic enzyme, is a top marker gene for nodule vasculature cells in scRNAseq data^38^, where GA accumulation is low (Fig. 1d). Localized post-transcriptional control of GA biosynthetic enzymes is an intriguing hypothesis for future studies. Assessing the role of localized transporters is also interesting albeit challenging to address as they are promiscuous and can transport molecules, such as nitrate, in addition to GA^57,58^.

The recruitment of GA in response to cytokinin treatment suggests a coherent feed-forward loop with a path both through and parallel to *NIN* underlying nodule organogenesis (Fig. 4). The loss of GA patterning in *noot* and *lsh* mutants is striking and sheds light on their roles as identity regulators, a key component which may be directing and maintaining hormone signaling in mature nodules as a mechanism contributing to their identity roles (Fig. 5, Fig. S8). Consistent with this, recent ChIP data for targets of LSH1 include a set of GA biosynthesis genes, including MtGA20ox1 (Medtr1g102070), one of the highest expressed GA20ox genes during nodulation, which provides a potentially direct link between LSH signalling and GA regulation^18,19^. The rescue in nodule size by GA treatment in *lsh1lsh2* mutants further supports that the perturbed GA pathway directly affects nodule size maintenance. The dependence on cytokinin for GA accumulation helps explain the strong differentiator that GA demonstrates between a nodule meristem and a lateral root meristem. We can now state that both the accumulation of cytokinin and GA helps differentiate nodulation from lateral root development and perhaps it is the accumulation of these hormones that dictates nodule identity over lateral root identity.

We demonstrate that targeted manipulation of GA levels can be leveraged to regulate nodule size, a result that is consistent with GA’s general role in growth and development (Fig. 3e,f), and a discovery that may be of interest in efforts to improve the efficiency of nitrogen-fixation. We speculate that there exists spatially resolved accumulations and depletions of GA that explain the concurrent roles of GA and DELLA in the rhizobium symbiosis. Taken together, our results demonstrate the importance of local GA accumulation for nodule primordia development through maturation and highlight how the ability to alter plant hormones in specific tissues potentiates novel aspects of plant development in diverse plant species.

## MATERIALS & METHODS

### Plant materials

The nlsGPS2 was genetically integrated into the R108 background of *M. truncatula* by the crop transformation team at the John Innes Centre. The NF19087 and NF13294 insertional mutants were ordered from the Oklahoma State University Medicago stock center. Genotyping primers and information can be found in supplemental Table S1.

### Cloning

The full length CDS for nlsGPS2 with domesticated BsaI and BpiI sites was synthesized (GeneWiz) into a MoClo compatible level 0 vector with SC overhangs^59^. This fragment was used with a compatible *L. japonicus UBIQUITIN* promoter (1.118 kb upstream) in a level 0 with PU overhangs and a t35S terminator with ST overhangs. Fragments were assembled into a level 1 vector, followed by assembly with a BASTA resistance cassette and an insulator fragment between the BASTA cassette and sensor following the protocol described in Engler et al. 2014^59^.

For misexpression of *MtGAOL1*, a full length CDS with domesticated for BsaI and BpiI sites was synthesized (GeneWiz) and amplified with C overhangs. This fragment was combined with a domesticated YPET with S overhangs, a tNOS terminator with ST overhangs and (1) a PU-flanked *MtFLOT4* promoter (2.0 kb upstream), (2) a PU-flanked AtCO2 promoter (647 bp upstream AT1G62500) or (3) a synthetic NIN promoter containing the 897 bp containing CE motif flanked by P sites and the 4.5 kb upstream region flanked by U sites by GoldenGate reaction into a level 1 vector. For over-expression of GA biosynthesis, the same NIN synthetic promoter described in the former was combined with a domesticated CDS of MtGA20ox1 flanked by S overhangs, a domesticated CDS of P2A flanked by C3 overhangs and a CDS of *MtGA3ox1* flanked by C4 overhangs, with a 35s terminator flanked by ST overhangs. These were combined into a level 1 vector by GoldenGate reaction. The promoters of *MtGA20ox1* (Medtr1g102070) and *MtGA3ox1* (Medtr2g102570) were synthesized (Genewiz) flanked by PU overhangs and combined with an NLS-3xVenus cassette flanked by SC overhangs and a 35s terminator flanked by ST overhangs. These were combined into a level 1 vector by GoldenGate reaction. For induction of spontaneous nodules construct, the *A. thaliana* UBQ10 promoter (1.5 kb upstream) flanked by PU sites was combined with the truncated CDS of MtCCaMKΔ1-311 flanked by SC sites and the *P. sativum* rubisco terminator flanked by ST sites into a level 1 construct. All hairy root level 1 constructs described were assembled into level 2 binary destination vectors that included a selection cassette in position 1 containing a ubiquitously-expressed (*A. thaliana* UBQ10) upstream an AtPIP2A coding sequence (AT3G53420) tagged with mScarlet-i. The control vector included the position 1 casette described in the former and an insulator cassette in position 2.

### *Agrobacterium rhizogenes* hairy root transformation

With the exception of the experiments in Fig. S4 using R108 seeds, *M. truncatula* A17 seeds were sterilized with 12% hypochlorite solution for 3 min, rinsed several times with sterile water, and washed for 1-4 hours at room temperature in sterile water containing 5 µM nystatin. Seeds were incubated for 3-6 days on 1% agar plates containing nitrogen-free BNM media^60^ containing 0.1µM GA4 and 5 µM nystatin. Seedlings were germinated for 24 hours at 24C in the dark followed by plating vertically on M-media (0.8 mM KNO_3_ · 3.0 mM MgSO_4_ · 0.9 mM KCl · 35 µM KH_2_PO_4_ · 1.2 mM Ca(NO3)_2_ · 30 µM MnCl_2_ · 24 µM H_3_BO_3_ · 9.2 µM ZnSO_4_ · 11.65 nM Na_2_MoO_4_ · 0.52 µM CuSO_4_ · 40 µM Glycine · 280 µM Myoinsitol · 4 µM Nicotinic acid · 0.5 µM Pyridoxine · 0.3 µM Thiamine · 1% Agarose) containing a layer of sterile filter paper. Seeds were sealed with micropore tape and roots were shaded from the light with black plastic. At 2-5 days old, seedlings were sliced at the root-hypocotyl junction and swirled in a prepared mix of *A. rhizogenes* (AR1193e). Prepared mix consisted of cultures grown to OD∼1.2, spun down and washed twice with sterile water, and resuspended in sterile water to an OD of 1.0 and a final concentration of 200µM Acetosyringone. Transformed seedlings were layered with sterile, wet filter paper and plates were then sealed parafilm. Seedlings were grown at 20C for one week in low light (∼40 µmol/m^2^). After one week, roots were transferred to M-media (see above) containing 250µg/ml Cefotaxime. For experiments using constructs with tissue-specific constitutive expression of *MtGAOL* or their respective controls, plates were supplemented with 1µM GA3 to overcome transformation defects from overexpression of GA catabolic enzymes. After a further two weeks, transformation was screened by checking for red fluorescent root systems. Untransformed roots were excised and removed before further experimentation.

### Inoculation of *M. truncatula* whole and composite plants in sand mix

A mix of 1:1:1 sand : terragreen : vermiculite was prepared in 24T pots. Sand was soaked and mixed with filtered water containing *S. meliloti* Sm2011 containing *nifH::*GUS at an OD = 0.1 Sm2011 was previously grown in minimal media at 28°C to an OD = 0.8 – 1.0. Sand mixture was spread evenly into pots. Plants were grown in walk-in chambers (22°C 16h light, 20°C 8h dark) for 2-4 weeks as specified in each experiment, with watering with liquid Fahräeus^61^ medium the first two weeks.

### Inoculation of *M. truncatul*a on agar-based plates

*M. truncatula* seeds were sterilized with 12% hypochlorite solution for 3 min, rinsed several times with sterile water, and washed for 1-4 hours at room temperature in sterile water containing 5 µM Nystatin. Seeds were incubated at 4°C for 3-5 days on BNM media^60^ containing 0.1µM GA_4_ and 5 µM nystatin. Seedlings were germinated for 24 hours at 24°C in the dark followed by plating vertically on BNM-media supplemented with 0.1µM aminoethoxyvinylglycine (AVG) with a piece of sterile filter paper between the media and the seedlings. Plates were sealed with micropore tape and roots were shaded from the dark with black plastic and grown in standard long day conditions (24°C 16h light, 22°C 8h dark). After two days, plants were spray inoculated with *S. meliloti* Sm2011 containing *nifH*::GUS (OD = 0.025) diluted in liquid Fahräeus medium^61^. The position of the root susceptible zone was marked with a sterile needle at the time of infection.

### Imaging nlsGPS2 in whole mount roots

For experiments at 4 or 5 days after infection, roots were placed on 10cm round petri dishes containing solidified BNM media (without filter paper). The areas around the marked susceptibility zone, or around lateral roots for lateral root imaging, were covered with 4-5% agarose cooled but not yet solidified. After solidification, plates were flooded with liquid BNM media. Imaging was carried out with a 25x water-dipping lens on an SP8-FLiman (Leica) with HyD detectors. For FRET measurement, the same settings described in Rizza et al. 2017^40^ were applied. Briefly, a 440nm laser was used for excitement and emission was collected at CFP (425-475 nm) and YFP (525-575 nm) ranges. For the YFP control channel, a 514 nm laser was used for excitement and YFP (525-575 nm) emission was used.

### Imaging nlsGPS2 in nodule slices/fluorescence microscopy

Nodules were embedded in 4-5% agarose and sliced in 100um sections using a vibratome (Leica). Samples were immediately imaged on slides using a 20x air objective using the excitation and emission settings described above. All images were acquired on the SP8-Fliman with HyD detectors or SP8-iPhox with HyD detectors (inverted). For images acquired on the SP8-iPhox, emission ratios are divided by 3 to be equivalent to those collected on the SP8-Fliman. For embedding nodule primordia, slices were mounted in 0.5% calcofluor white in sterile water to observe cell walls.

### Confocal Image Analysis and emission ratio quantification

For demonstrative images, Fiji was used to generate sum stacks. Display images (i.e. those not used for quantification) of YFP, CFP, and FRET channels, the channel brightness has been adjusted and background was removed by assessing the brightness histogram of the background (outside of the root/nodule sample). The FRETENATOR2 plugin was used in FIJI for all emission ratio analysis^62^. Briefly, nuclei segmentation was selected using the YFP control channel. The emission ratio was calculated per nuclei by evaluation of FRET/CFP channel. Saturated nuclei, non-segmented nuclei, or nuclei too dim in donor CFP channel (sum CFP brightness must be 5% higher than background) are excluded from analysis. For determining nodule center, bright field images were observed for x and y position (blinded from emission ratio outcome).

### Transmission Electron Microscopy

TEM was carried out on 4 wpi nodules inoculated in sand mix with Sm2011. Protocol is described in Schiessl et al. 2023^19^.

### Image analysis for assessing nodule length

For measuring nodule length from base to tip, images were taken and measured on a VHX-7000 (Keyence) or on a DS-7000 Flatbed scanner (EPSON). For images shown from the flatbed scanner, the image background was removed in Fiji (50 pixel rolling ball radius).

### Plotting and statistics

R software was used for generating all plots and statistics. For statistical tests, data was tested for normality distribution by Shapiro-Wilkes test. For normally distributed data, an F-test was used to assess variance. If groups had equal variance, a Student’s t-test was used to evaluate samples. If the F-test had a p-value <0.05, a Welch’s two sample t-test was used. For non-normally distributed data, a Wilcoxon-Rank Sum test was used, unless there was a sample size of N > 20, in which case a Welch’s two sample test was used.

## SUPPLEMENTAL INFORMATION

**Fig. S1.**
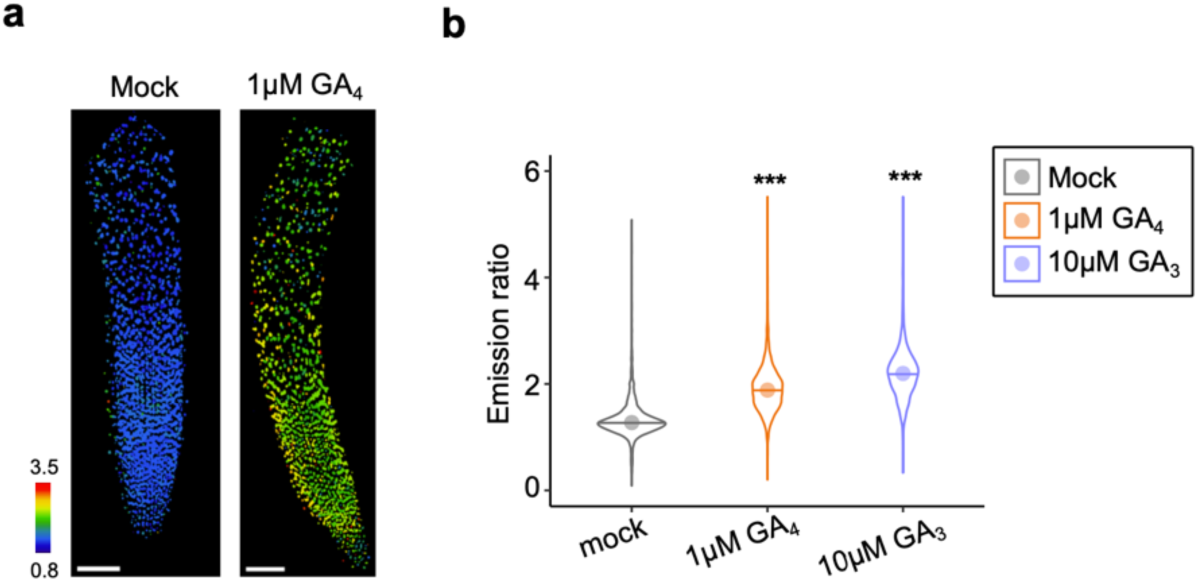
Response of genetically-encoded GA sensor in *M. truncatula*. **(a)** Emission ratio of *M. truncatula* roots containing genetically-encoded nlsGPS2 inoculated for 20 min with mock or 1µM GA_4_. Bar = 100 µm. (**b**) Emission ratio distribution of nuclei from roots treated for 20 min with mock, 1µM GA_4_ or 10µM GA_3_. N≥3. Welch’s t-test ***p-value < 0.001. Median values: mock = 1.274, 1 µM GA_4_ = 1.885, 1µM GA_3_ = 2.197.

**Fig. S2.**
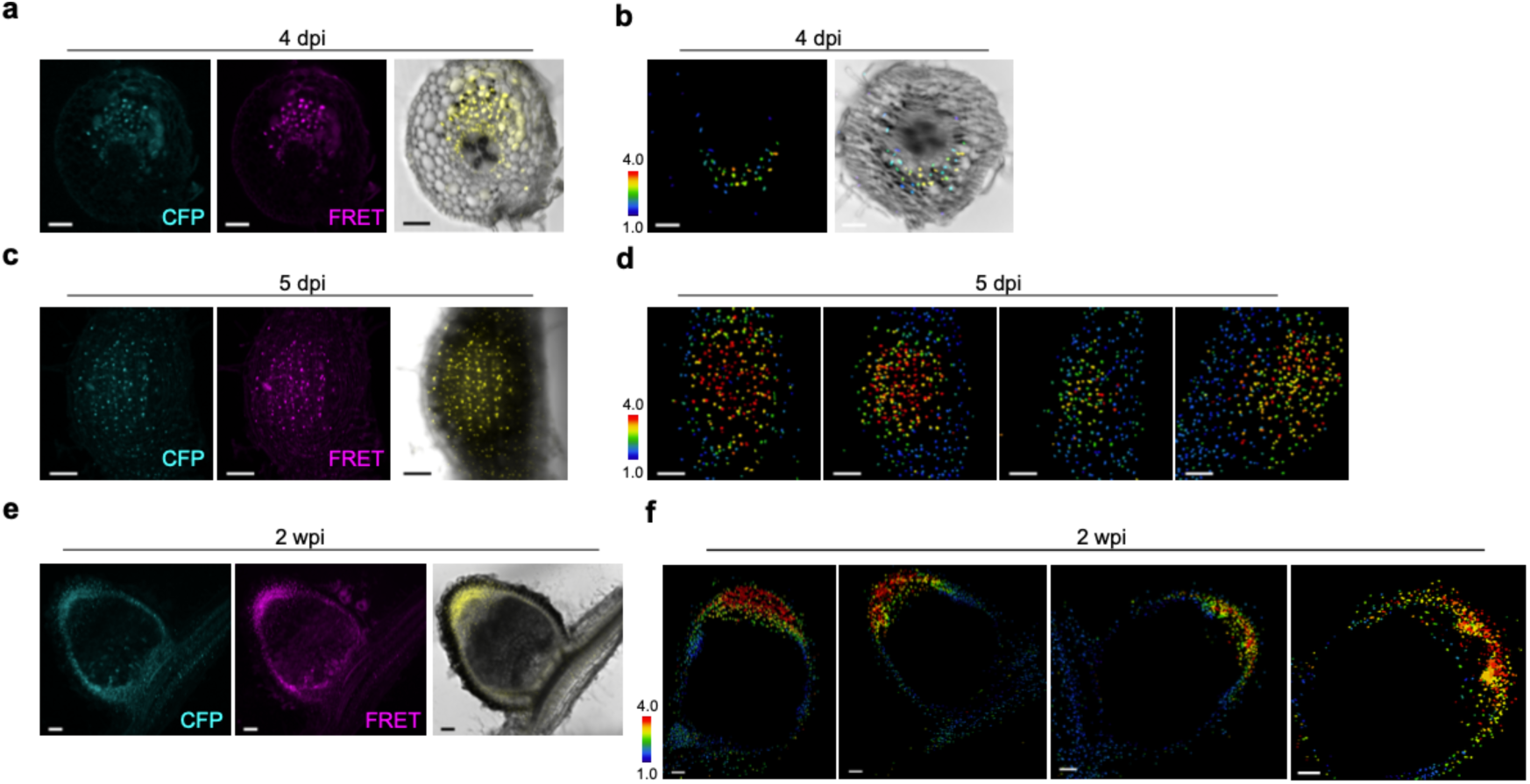
Additional examples of GA accumulation in nodule development. **(a)** Sum projection of CFP, FRET and brightfield/YFP channels from sample in Fig. 1b. **(b)** Emission ratio and YFP/brightfield overlay of additional example of 4 dpi nodule primordia embedded in 4.5% agarose and sliced in 100 µm sections. **(c)** Sum projection of CFP, FRET and brightfield/YFP channels from sample in main Fig. 1c. **(d)** Additional examples of emission ratio of 5 dpi nodule primordia, wholemount. **(e)** Sum projection of CFP, FRET and brightfield/YFP channels from sample in main Fig. 1d. **(f)** Additional examples of emission ratio of 2 wpi nodule primordia, embedded in 4.5% agarose and sliced in 100 µm sections.

**Fig. S3.**
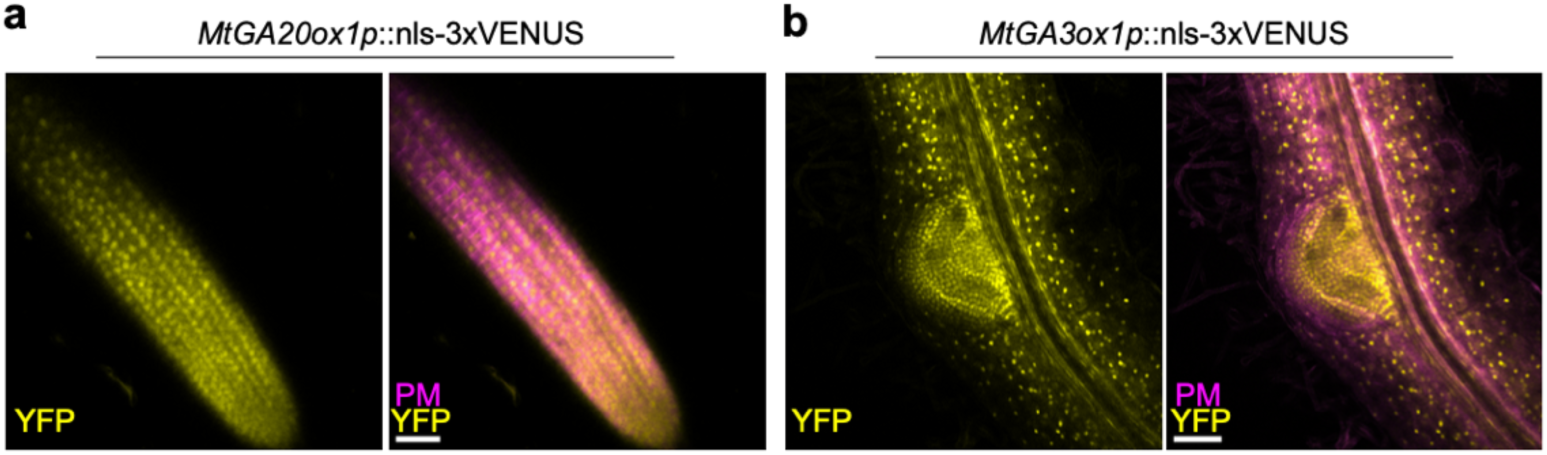
Expression of GA biosynthesis genes in *A. rhizogenes*-transformed *M. truncatula* roots. **(a)** Expression of *MtGA20ox1(3kb)p::nls-3xVenus* in the primary root, N=10. **(b)** Expression of *MtGA3ox1p(3kb)p::*nls-3xVenus in the primary root and lateral root primordia, N=10. PM = ubiquitous plasma-membrane marker tagged with mScarlet (*AtUBQp::*AtPIP2A-mScarleti) present in vector used for selection. Bar = 50µm.

**Fig. S4.**
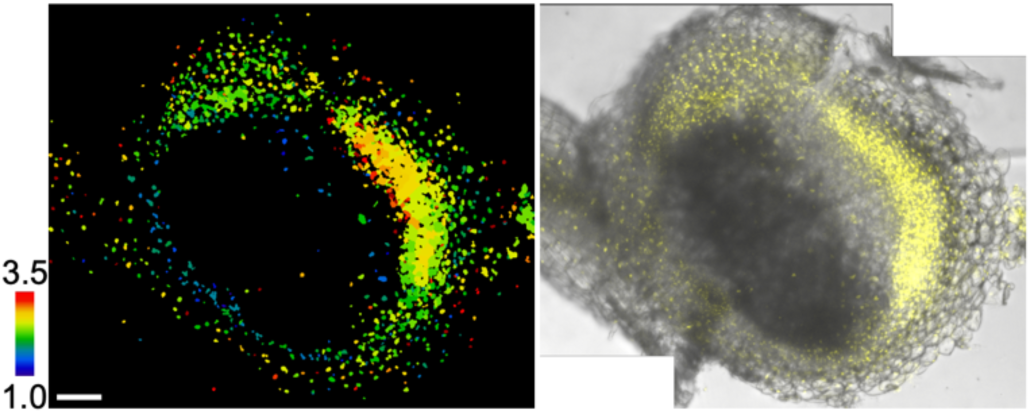
GA accumulation in *A. rhizogenes*-transformed spontaneous nodules. Emission ratio image (left) and YFP/Brightfield overlay of *LjUBQp*::nlsGPS2 *M. truncatula* transformed with *AtUBQ10::*MtCCaMKΔ1-311. Spontaneous nodules form after 5 weeks. N=3.

**Fig. S5.**
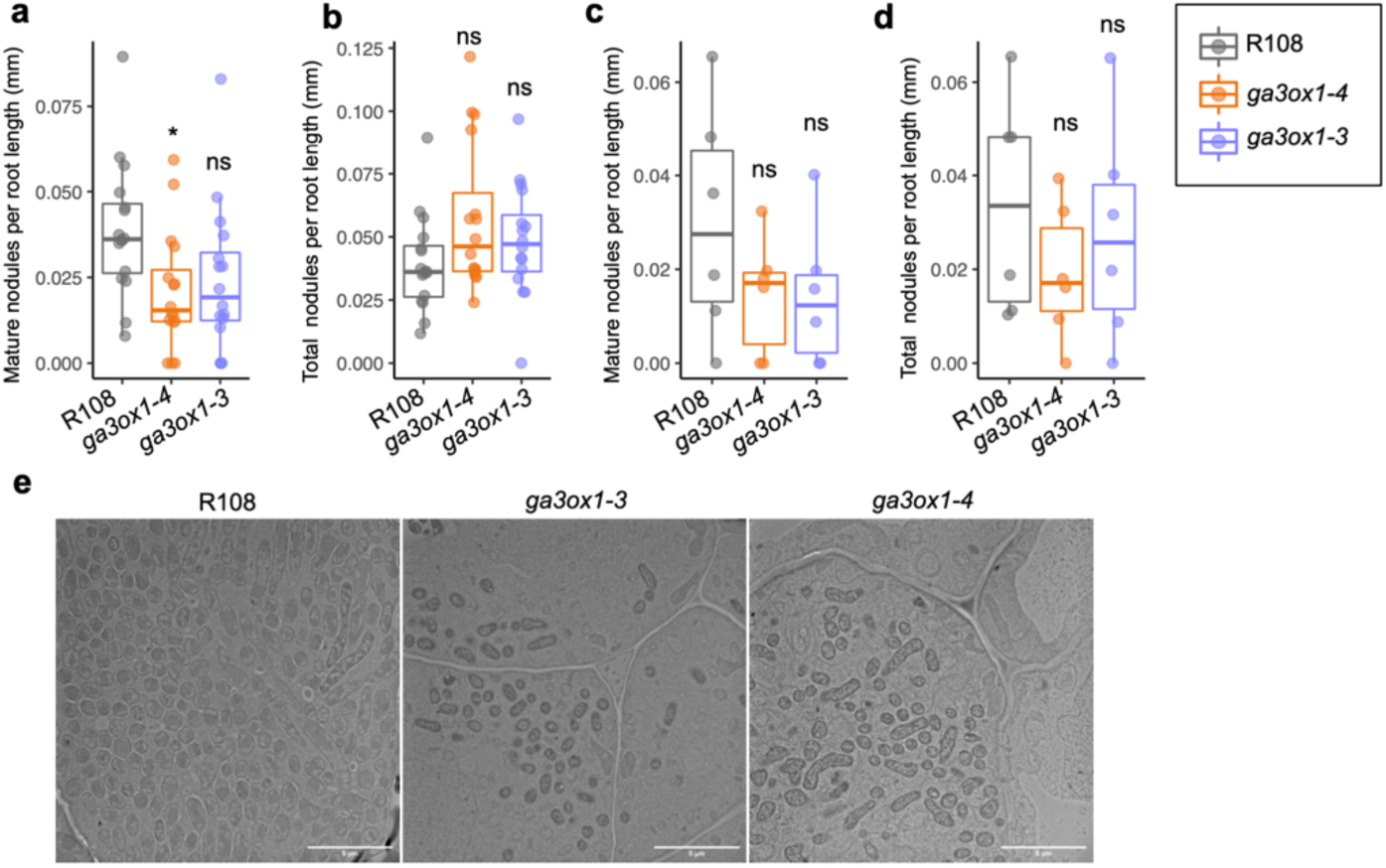
Further *ga3ox1* mutant phenotypes. Quantification of **(a)** mature (pink) nodules and **(b)** total nodules per root length (mm) for R108, *ga3ox1-3* and *ga3ox1-4* inoculated with *S. meliloti* Sm2011 on plates for 14 dpi. N=16, Student’s t-test p-value * < 0.05. Median values in (a): R108 = 0.036, *ga3ox1-4* = 015, *ga3ox1-3* = 0.019. Median values in (b): R108 = 0.036, *ga3ox1-4* = 0.0462, *ga3ox1-3* = 0.0471. Quantification of **(c)** mature (pink) and **(d)** total nodules per root length (mm) for R108, *ga3ox1-3* and *ga3ox1-4* inoculated with Sm2011 in sand mix for 14 dpi. Student’s t-test, not significant. Median values in (c): R108 = 0.027, *ga3ox1-4* = 0.017, *ga3ox1-3* = 0.012. Median values in (d): R108 = 0.03, *ga3ox1-4* = 0.017, *ga3ox1-3* = 0.026. (**e**) TEM images 2600x of apical infection zone of R108, *ga3ox1-3* and *ga3ox1-4.* Bar = 5µm.

**Fig. S6.**
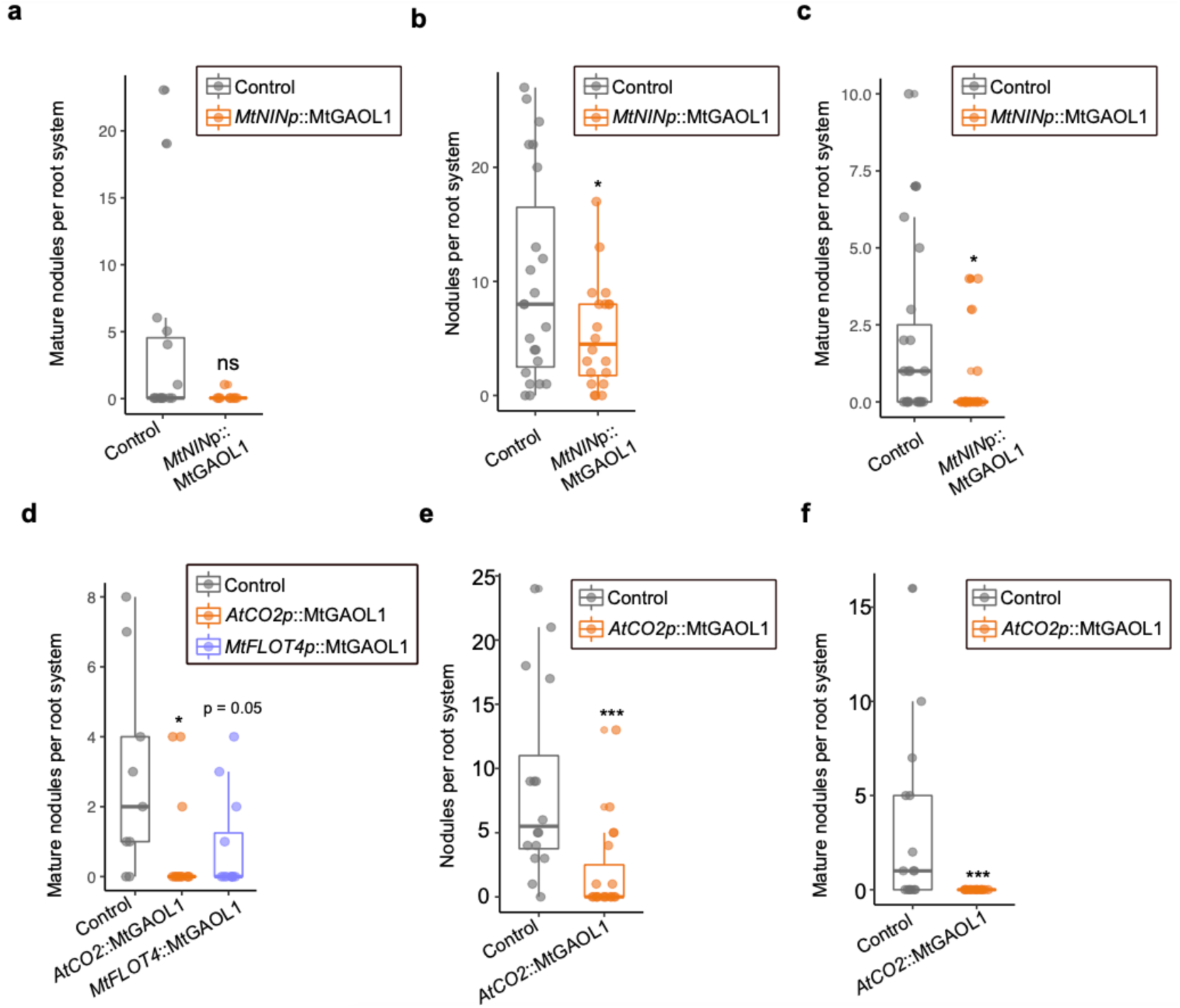
Supporting information for the function of GA in developing nodule primordia cortical cells. (a) Total mature (pink) nodules per root systems from replicate displayed in Fig. 3f. Median values: control = 0, MtNINp::GAOL1 = 0. Wilcox rank sum test, not significant. N≥10. (b) Total nodules per root system of second replicate of data in Fig. 3f. Welch’s t-test *p-value < 0.05, median values: control = 8, GAOL1 = 5. N≥ 21. (c) Mature (pink) nodules per root system of second replicate of experiment in Fig. 3f. Welch’s t-test *p-value < 0.05. Median values: control = 1, MtNINp::GAOL1 = 0. N≥ 21 (d) Total mature (pink) nodules per root systems from data in Fig. 3g. Wilcox test *p-value < 0.05, p-value for MtFLOT4p::GAOL1 = 0.0499. Median values: control = 2, *AtCO2p::*GAOL1 = 0, *MtFLOT4p*::GAOL1 = 0. N≥10. **(e)** Total nodules per root system in second replicate of data in Fig. 3g. Wilcoxon rank sum test ***p-value < 0.001. Median values: control = 5.5, *AtCO2p::*GAOL1 = 0. N≥16. **(f)** Mature (pink) nodules of second replicate of data in Fig. 3g. Wilcoxon rank sum test ***p-value <0.001. Median values: control = 1, *AtCO2p::*GAOL1 = 0. N≥16.

**Fig. S7.**
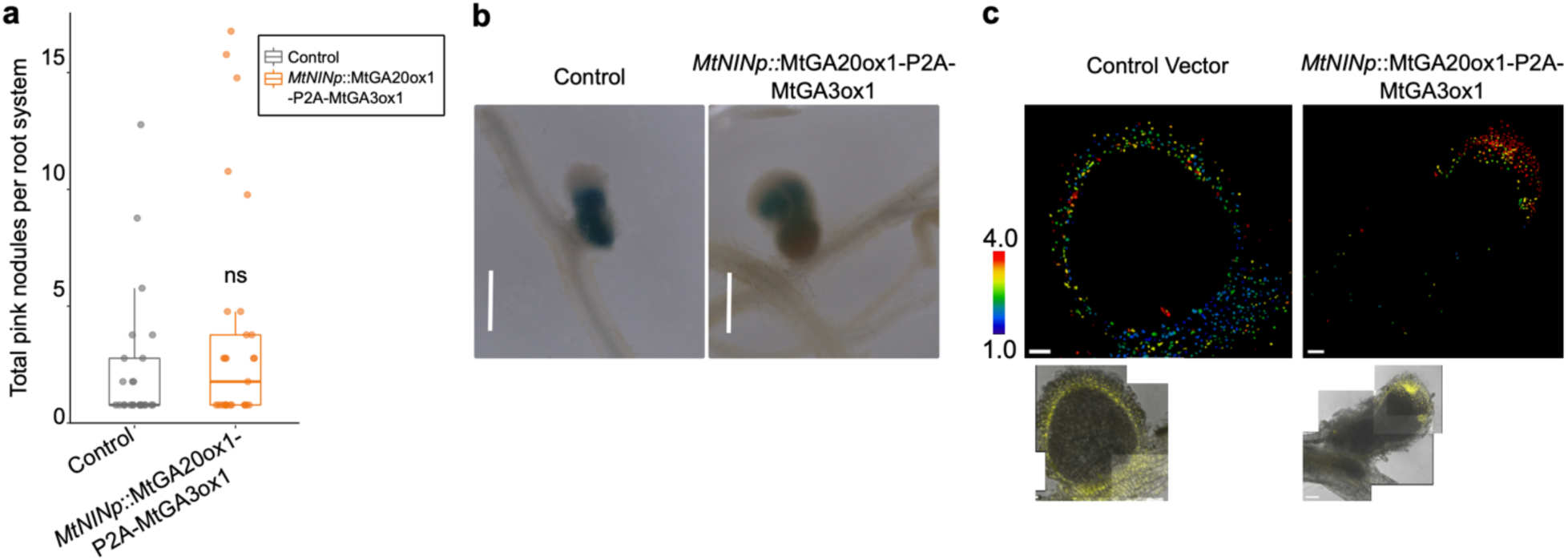
Supporting information for the function of GA in nodule size. **(a)** Total mature nodules per root system in *A. rhizogenes* roots transformed with red plasma-membrane casette control (*AtUBQp*::PM-mScarleti) or *MtNINp::*MtGA20ox1-P2A-MtGA3ox, plasma-membrane cassette. Welch’s t-test, not significant. Median values: control = 0, *MtNINp*::MTGA20ox1-P2A-MtGA3ox1 = 1. N≥25**. (b)** Representative images of nodules from control and *MtNINp::*MtGA20ox1-P2A-MtGA3ox1-transformed roots. Bar = 1000µm. Blue is indicative of GUS staining for Sm2011 *nifH::*GUS, which express GUS under a symbiotic-regulated operon. **(c)** Emission ratio (top panel) of *LjUBQp::*nlsGPS2 *M. truncatula* roots transformed with control or *MtNINp::*MtGA20ox1-P2A-MtGA3ox1 and YFP, brightfield (bottom panel), N=3.

**Fig.S8.**
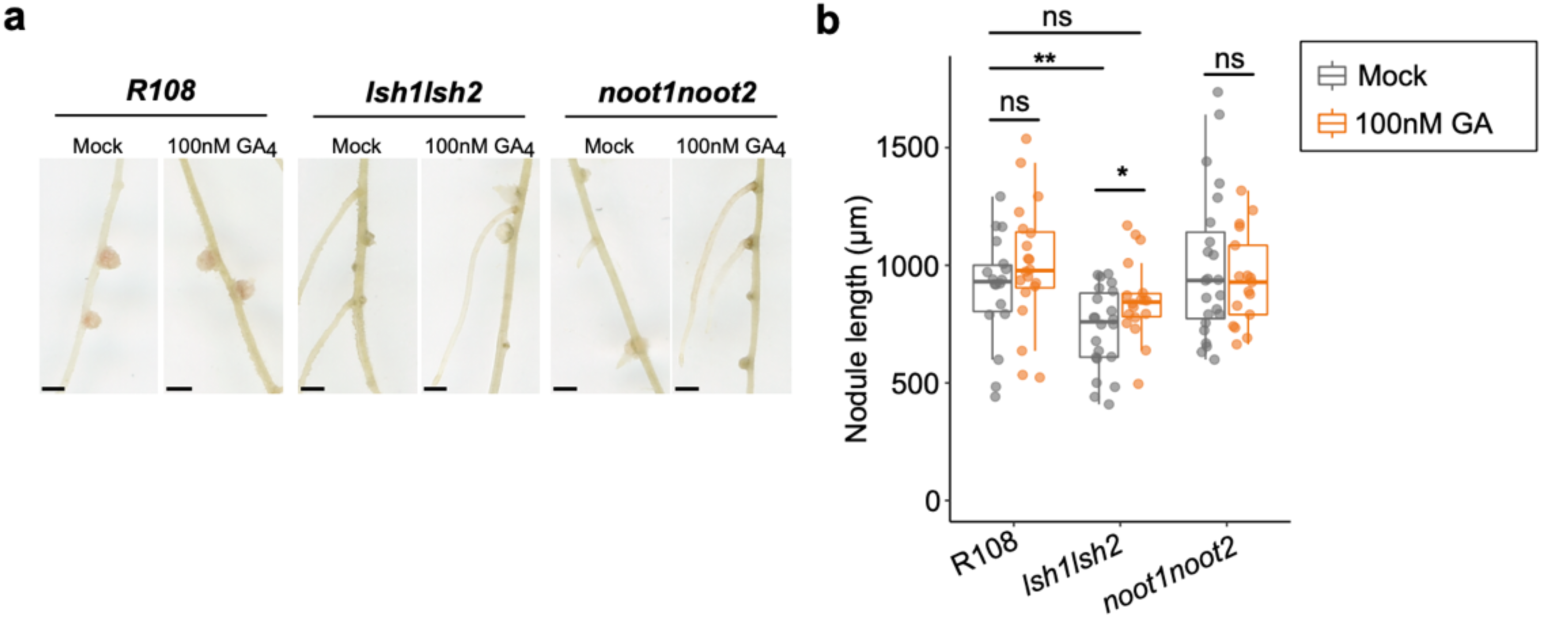
GA treatment after nodule initiation of wild-type and organogenesis mutations. **(a)** Representative scans of R108, *lsh1lsh2* and *noot1noot2* at 14 dpi treated at 4 dpi with mock or 100nM GA_4_. Bar = 1.0 mm. **(b)** Quantification of nodule length at 14 dpi for plants treated with mock or 100nM GA_4_. Median values R108: mock = 930.45, GA_4_ = 977.49; median values *lsh1lsh2*: mock = 759.52, *l*GA_4_ = 843.59; median values *noot1noot2:* mock = 935.12, GA_4_ = 928.57; Student’s t-test *p-value <0.05, **p-value < 0.01. N≥17. Data shown from three biological replicates.

**Fig. S9.**
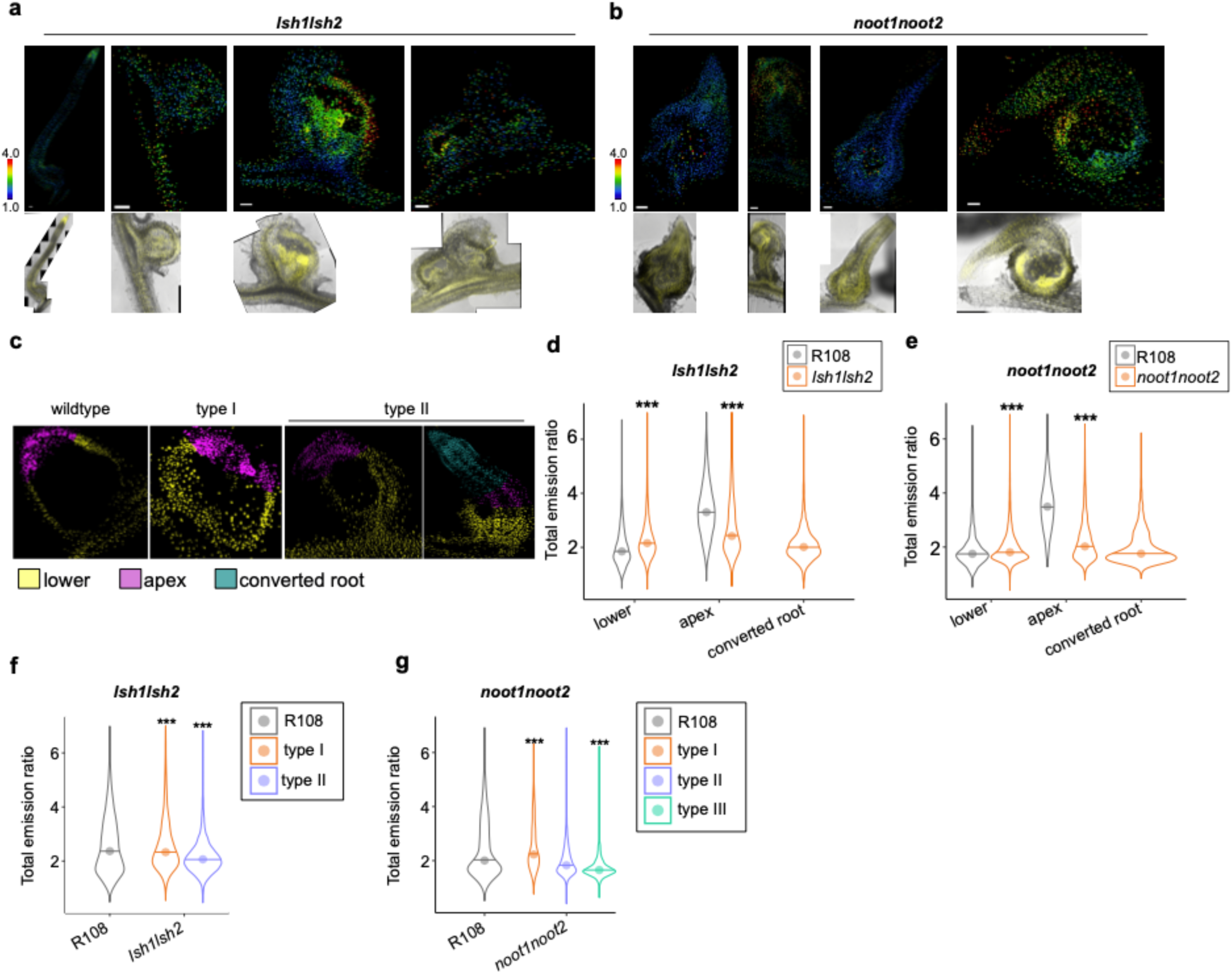
Nodule organogenesis regulators function in GA accumulation in mature nodules. **(a)** Example emission ratios from *lsh1lsh2* mutants (top panel) and their respective YFP/brightfield control channels from 2wpi nodules. **(b)** Example emission ratios from *lsh1lsh2* mutants (top panel) and their respective YFP/brightfield control channels from 2 wpi nodules. **(c)** Examples of region determination using FRETENATOR ROI labeling map. **(d)** Emission ratio distribution of nuclei by region in lower nodule/root, nodule apex, or converted lateral root portion (mutant only) between nlsGPS2 and *lsh1lsh2*; Welch’s T-test, **p-value <0.01,***p-value <0.001. Median values R108: lower = 1.852, apex = 3.3115; median values *lsh1lsh2:* lower = 2.159, apex = 2.441, converted root = 2.012. N≥15. **(e)** Emission ratio distribution of nuclei by region in lower nodule/root, nodule apex, or converted lateral root portion (mutant only) between nlsGPS2 and *noot1noot2.* Welch’s T-test, ***p-value <0.001. Median values R108: lower = 1.749, apex = 3.501. Median values *noot1noot2:* lower = 1.806, apex = 2.026, converted root = 1.7545. N≥15. Emission ratio plotted in (d-e) are from the same dataset presented in main Figure 5**.(f)** Emission ratio distribution of organ types in R108 and *lsh1lsh2* mutants. Welch’s T-test, ***p-value <0.001. Median values: R108 = 2.373, type I = 2.34, type II = 2.065. N≥15. (g). Emission ratio distribution of organ types in R108 and *noot1noot2* mutants. Welch’s T-test, ***p-value <0.001. Median values: R108 = 2.0045, type I = 2.231, type II = 1.831, type III = 1.653.N≥15. Data shown are from three biological replicates.

## ACKNOWLEDGMENTS

We are grateful to Wendy Harwood and her crop transformation at the John Innes Centre for producing the nlsGPS2 stable lines. We thank Kim Findlay and Elaine Barclay at the John Innes Centre for their TEM service. We thank Aleksander Gavrin and Defeng Shen for their *A. rhizogenes* transformation technical suggestions. We are also grateful to Tonni Andersen, Wouter Kohlen, James Rowe, Edwin Jarratt Barnham, Chenwu Liu and all of the Jones team for their scientific discussions that improved this manuscript. This work was supported by H2020-IF (844398) to CD, the European Molecular Biology Organization (ALTF-111-2019) to CD, the Gatsby Charitable trust (GAT3395) to AMJ and GEDO, and the Bill and Melinda Gates Foundation and the UK Foreign, Commonwealth and Development Office (OPP1028264) through Engineering the Nitrogen Symbiosis for Africa (ENSA) project to GEDO.

## CONTRIBUTIONS

CD, AMJ and GEDO conceived and designed the experiments. CD, AR, NRM, and FDS carried out the experimental work. CD, KS and JW contributed unpublished plant materials. CD, AMJ, GEDO, AR, KS, NRM, FDS contributed intellectually to the manuscript. CD, AMJ, and GEDO wrote the manuscript.

## REFERENCES

1. Oldroyd, G. & Leyser, O. A plant’s diet, surviving in a variable nutrient environment. Science 1, 6–21 (2020).

2. Jhu, M.-Y. & Oldroyd, G. E. D. Dancing to a different tune, can we switch from chemical to biological nitrogen fixation for sustainable food security? PLOS Biol. 21, e3001982 (2023).

3. Gonzalez-Rizzo, S., Crespi, M. & Frugier, F. The Medicago truncatula CRE1 Cytokinin Receptor Regulates Lateral Root Development and Early Symbiotic Interaction with Sinorhizobium meliloti. Plant Cell 18, 2680–2693 (2006).

4. Murray, J. D. et al. A Cytokinin Perception Mutant Colonized by Rhizobium in the Absence of Nodule Organogenesis. Science 315, 101–104 (2007).

5. Tirichine, L. et al. A Gain-of-Function Mutation in a Cytokinin Receptor Triggers Spontaneous Root Nodule Organogenesis. Science 315, 104–107 (2007).

6. Reid, D. et al. Cytokinin Biosynthesis Promotes Cortical Cell Responses during Nodule Development. Plant Physiol 175, 361–375 (2017).

7. Marsh, J. F. et al. Medicago truncatula NIN Is Essential for Rhizobial-Independent Nodule Organogenesis Induced by Autoactive Calcium/Calmodulin-Dependent Protein Kinase. Plant Physiol. 144, 324–335 (2007).

8. Heckmann, A. B. et al. Cytokinin Induction of Root Nodule Primordia in Lotus japonicus Is Regulated by a Mechanism Operating in the Root Cortex. Mol. Plant-Microbe Interact. 24, 1385–1395 (2011).

9. Plet, J. et al. MtCRE1-dependent cytokinin signaling integrates bacterial and plant cues to coordinate symbiotic nodule organogenesis in Medicago truncatula. Plant J. 65, 622–633 (2011).

10. Gauthier-Coles, C., White, R. G. & Mathesius, U. Nodulating Legumes Are Distinguished by a Sensitivity to Cytokinin in the Root Cortex Leading to Pseudonodule Development. Front. Plant Sci. 9, 1901 (2019).

11. Liu, J. et al. NIN is essential for development of symbiosomes, suppression of defence and premature senescence in Medicago truncatula nodules. N. Phytol. 230, 290–303 (2021).

12. Feng, J., Lee, T., Schiessl, K. & Oldroyd, G. E. D. Processing of NODULE INCEPTION controls the transition to nitrogen fixation in root nodules. Science 374, 629–632 (2021).

13. Yoro, E. et al. A Positive Regulator of Nodule Organogenesis, NODULE INCEPTION, Acts as a Negative Regulator of Rhizobial Infection in Lotus japonicus. Plant Physiol. 165, 747–758 (2014).

14. Liu, J. et al. A Remote cis-Regulatory Region Is Required for NIN Expression in the Pericycle to Initiate Nodule Primordium Formation in Medicago truncatula. Plant Cell 31, 68–83 (2019).

15. Couzigou, J.-M. et al. NODULE ROOT and COCHLEATA Maintain Nodule Development and Are Legume Orthologs of Arabidopsis BLADE-ON-PETIOLE Genes. Plant Cell 24, 4498–4510 (2012).

16. Magne, K. et al. MtNODULE ROOT1 and MtNODULE ROOT2 are essential for indeterminate nodule identity. Plant Physiol 178, pp.00610.2018 (2018).

17. Shen, D. et al. A Homeotic Mutation Changes Legume Nodule Ontogeny into Actinorhizal-Type Ontogeny. Plant Cell 32, 1868–1885 (2020).

18. Schiessl, K. et al. NODULE INCEPTION Recruits the Lateral Root Developmental Program for Symbiotic Nodule Organogenesis in Medicago truncatula. Curr Biol 29, 3657–3668.e5 (2019).

19. Schiessl, K. et al. Light sensitive short hypocotyl (LSH) confer symbiotic nodule identity in the legume Medicago truncatula. Biorxiv 2023.02.12.528179 (2023) doi:10.1101/2023.02.12.528179.

20. Soyano, T., Shimoda, Y., Kawaguchi, M. & Hayashi, M. A shared gene drives lateral root development and root nodule symbiosis pathways in Lotus. Science 366, 1021– 1023 (2019).

21. Laporte, P. et al. The CCAAT box-binding transcription factor NF-YA1 controls rhizobial infection. J. Exp. Bot. 65, 481–494 (2014).

22. Laloum, T. et al. Two CCAAT-box-binding transcription factors redundantly regulate early steps of the legume-rhizobia endosymbiosis. Plant J. 79, 757–768 (2014).

23. Xiao, T. T. et al. Fate map of Medicago truncatula root nodules. Development 141, 3517–3528 (2014).

24. Floss, D. S., Levy, J. G., Lévesque-Tremblay, V., Pumplin, N. & Harrison, M. J. DELLA proteins regulate arbuscule formation in arbuscular mycorrhizal symbiosis. Proc National Acad Sci 110, E5025–E5034 (2013).

25. Fonouni-Farde, C. et al. DELLA-mediated gibberellin signalling regulates Nod factor signalling and rhizobial infection. Nat Commun 7, 12636 (2016).

26. Jin, Y. et al. DELLA proteins are common components of symbiotic rhizobial and mycorrhizal signalling pathways. Nature Communications 7, 12433 (2016).

27. Maekawa, T. et al. Gibberellin controls the nodulation signaling pathway in Lotus japonicus. Plant J. 58, 183–194 (2009).

28. Fonouni-Farde, C. et al. DELLA1-Mediated Gibberellin Signaling Regulates Cytokinin-Dependent Symbiotic Nodulation. Plant Physiol 175, 1795–1806 (2017).

29. Ferguson, B. J., Ross, J. J. & Reid, J. B. Nodulation Phenotypes of Gibberellin and Brassinosteroid Mutants of Pea. Plant Physiol. 138, 2396–2405 (2005).

30. McAdam, E. L., Reid, J. B. & Foo, E. Gibberellins promote nodule organogenesis but inhibit the infection stages of nodulation. J Exp Bot 69, 2117–2130 (2018).

31. Benedito, V. A. et al. A gene expression atlas of the model legume Medicago truncatula. Plant J. 55, 504–513 (2008).

32. Limpens, E. et al. Cell- and Tissue-Specific Transcriptome Analyses of Medicago truncatula Root Nodules. PLoS ONE 8, e64377 (2013).

33. Roux, B. et al. An integrated analysis of plant and bacterial gene expression in symbiotic root nodules using laser-capture microdissection coupled to RNA sequencing. Plant J 77, 817–837 (2014).

34. Akamatsu, A. et al. Endogenous gibberellins affect root nodule symbiosis via transcriptional regulation of NODULE INCEPTION in Lotus japonicus. Plant J 105, 1507–1520 (2021).

35. Chu, X., Su, H., Hayashi, S., Gresshoff, P. M. & Ferguson, B. J. Spatiotemporal Changes in Gibberellin Content are Required for Soybean Nodulation. New Phytol (2021) doi:10.1111/nph.17902.

36. Gao, J. et al. NODULE INCEPTION activates gibberellin biosynthesis genes during rhizobial infection. New Phytol (2023) doi:10.1111/nph.18759.

37. Cervantes-Pérez, S. A. et al. Cell-specific pathways recruited for symbiotic nodulation in the Medicago truncatula legume. Mol Plant doi:10.1016/j.molp.2022.10.021.

38. Ye, Q. et al. Differentiation trajectories and biofunctions of symbiotic and un-symbiotic fate cells in root nodules of Medicago truncatula. Mol. Plant 15, 1852–1867 (2022).

39. Nett, R. S., Bender, K. S. & Peters, R. J. Production of the plant hormone gibberellin by rhizobia increases host legume nodule size. Isme J 16, 1809–1817 (2022).

40. Rizza, A., Walia, A., Lanquar, V., Frommer, W. B. & Jones, A. M. In vivo gibberellin gradients visualized in rapidly elongating tissues. Nat Plants 3, 803–813 (2017).

41. Rizza, A., et al. Differential biosynthesis and cellular permeability explain longitudinal gibberellin gradients in growing roots. Proc. Natl. Acad. Sci. 118, e1921960118 (2021).

42. Gleason, C. et al. Nodulation independent of rhizobia induced by a calcium-activated kinase lacking autoinhibition. Nature 441, 1149–1152 (2006).

43. Hedden, P. & Thomas, S. G. Annual Plant Reviews, Volume 49. (2022) doi:10.1002/9781119210436.

44. Barker, R. et al. Mapping sites of gibberellin biosynthesis in the Arabidopsis root tip. New Phytol 229, 1521–1534 (2021).

45. Rival, P. et al. Epidermal and cortical roles of NFP and DMI3 in coordinating early steps of nodulation in Medicago truncatula. Development 139, 3383–3391 (2012).

46. Jardinaud, M.-F. et al. A Laser Dissection-RNAseq Analysis Highlights the Activation of Cytokinin Pathways by Nod Factors in the Medicago truncatula Root Epidermis. Plant Physiol. 171, 2256–2276 (2016).

47. Liang, P. et al. Symbiotic root infections in Medicago truncatula require remorin-mediated receptor stabilization in membrane nanodomains. Proc National Acad Sci 115, 201721868 (2018).

48. Hedden, P. The current status of research on gibberellin biosynthesis. Plant Cell Physiol 61, pcaa092-(2020).

49. Rutten, P. J. & Poole, P. S. Chapter Nine Oxygen regulatory mechanisms of nitrogen fixation in rhizobia. Adv. Microb. Physiol. 75, 325–389 (2019).

50. Tsien, R. Y. THE GREEN FLUORESCENT PROTEIN. Biochemistry 67, 509–544 (1998).

51. Cowling, R. J. & Harberd, N. P. Gibberellins control Arabidopsis hypocotyl growth via regulation of cellular elongation. J. Exp. Bot. 50, 1351–1357 (1999).

52. Zhu, Y. et al. ELONGATED UPPERMOST INTERNODE Encodes a Cytochrome P450 Monooxygenase That Epoxidizes Gibberellins in a Novel Deactivation Reaction in Rice. Plant Cell 18, 442–456 (2006).

53. Boss, P. K. & Thomas, M. R. Association of dwarfism and floral induction with a grape ‘green revolution’ mutation. Nature 416, 847–850 (2002).

54. Hytönen, T., Elomaa, P., Moritz, T. & Junttila, O. Gibberellin mediates daylength-controlled differentiation of vegetative meristems in strawberry (Fragaria × ananassaDuch). BMC Plant Biol. 9, 18 (2009).

55. Tenreira, T. et al. A Specific Gibberellin 20-Oxidase Dictates the Flowering-Runnering Decision in Diploid Strawberry. Plant Cell 29, 2168–2182 (2017).

56. Kim, G.-B., Son, S.-U., Yu, H.-J. & Mun, J.-H. MtGA2ox10 encoding C20-GA2-oxidase regulates rhizobial infection and nodule development in Medicago truncatula. Scientific Reports 1–13 (2019).

57. Kanno, Y. et al. AtSWEET13 and AtSWEET14 regulate gibberellin-mediated physiological processes. Nat Commun 7, 13245 (2016).

58. Chiba, Y. et al. Identification of Arabidopsis thaliana NRT1/PTR FAMILY (NPF) proteins capable of transporting plant hormones. J. Plant Res. 128, 679–686 (2015).

59. Engler, C. et al. A Golden Gate Modular Cloning Toolbox for Plants. Acs Synth Biol 3, 839–843 (2014).

60. Ehrhardt, D. W., Atkinson, E. M. & Long, S. R. Depolarization of Alfalfa Root Hair Membrane Potential by Rhizobium meliloti Nod Factors. Science 256, 998–1000 (1992).

61. Boisson-Dernier, A. et al. Agrobacterium rhizogenes -Transformed Roots of Medicago truncatula for the Study of Nitrogen-Fixing and Endomycorrhizal Symbiotic Associations. Mol Plant-microbe Interactions 14, 695–700 (2001).

62. Rowe, J. H., Rizza, A. & Jones, A. M. Environmental Responses in Plants, Methods and Protocols. Methods Mol. Biol. 2494, 239–253 (2022).

